# Testing the plasticity-first theory of tumor evolution with spatiotemporal lineage tracing

**DOI:** 10.1101/2025.08.06.668639

**Authors:** Jinhua Yang, Liangzhen Hou, Xue Wang, Ningxia Zhang, Yifei Bian, Zhaolian Lu, Yanxu Chen, Duo Xie, Yongting Fang, Kun Wang, Ruijie Wan, Yujuan Jin, Yuan Chen, Xinlei Cai, Xiaojing Hu, Leo Tsz On Lee, Xinyuan Tong, Xionglei He, Zheng Hu, Hongbin Ji

## Abstract

The plasticity-first hypothesis posits that phenotypic plasticity emerges first in novel environments, followed by (epi)genetic changes that stabilize adaptive phenotypes. Whether tumor evolution follows this model, and how non-genetic mechanisms sustain long-term therapeutic resistance, remain unclear. Here, we develop eTRACER, an endogenously targeted, high-resolution and spatially-resolved CRISPR-Cas9 lineage-tracing system, to study tumor adaptation during longitudinal CD8^+^ T cell-mediated immunotherapy. Integrated clonal and single-cell transcriptomic analyses show that tumor immune evasion is governed by clonal state composition rather than clonal identity, arguing against selection from pre-existing genetic heterogeneity. Spatiotemporal lineage tracing reveals highly reversible state transitions prior to therapy, followed by consolidated epithelial-mesenchymal transition (EMT) during therapy. Single-cell multiomics identifies AP-1 as a key regulator of EMT stabilization and its inhibition enhances tumor sensitivity to immune killing. Together, these findings show that tumor plasticity precedes stable immune evasion, revealing an AP-1-centered mechanism underlying plasticity-first tumor evolution.

## Introduction

Tumor cells frequently evolve resistance during drug treatment, ultimately leading to therapeutic failure and patient death. Classical theory posits that drug resistance evolves through a Darwinian selective process, whereby therapeutic pressure acts on pre-existing clonal heterogeneity to drive the emergence of resistant populations^1–3^. However, substantial evidence highlights the critical role of non-genetic mechanisms in drug resistance^4–7^, where tumor cells harness the capacity of lineage plasticity to acquire an adaptive trait. The plasticity-first hypothesis^6,8–11^ proposes that environmentally induced phenotypic plasticity emerges first in novel environments, followed by (epi)genetic changes that stabilize these adaptive phenotypes, rendering them heritable across generations. Therefore, the plasticity-first model reconciles genetic and non-genetic theory of adaptation, highlighting that non-genetic mechanisms promote immediate survival upon microenvironmental changes while (epi)genetic mechanisms promote long-term adaptation. Although the plasticity-first model has also been posited to underlie the evolution of cancer drug resistance^10,12^ and tumorigenesis in general^13^, direct evidence from lineage tracing studies is still lacking. In the context of immunotherapy, such as adoptive cell transfer^14^, whether tumor cells exploit a plasticity-first strategy to evade immune surveillance remains elusive^15,16^.

Lineage barcoding combined with single-cell multiomic technologies represents a powerful approach for resolving cellular dynamics and fate-determining mechanisms^17,18^. To record cell lineage information using gene editing technologies such as CRISPR/Cas9^19–27^, base editors^28–30^, or prime editors^31,32^, current lineage barcoding approaches typically engineer exogenous target arrays as evolving barcodes. A fundamental limitation of this strategy is that the high randomness of integration in the host genome leads to dramatic fluctuation in target expression and editing efficiency across clones and cell types^33^. Such variability introduces substantial bias in lineage tree reconstruction, potentially skewing downstream evolutionary and functional analyses. Furthermore, current CRISPR/Cas9 lineage tracers often employ tandem target arrays that are susceptible to large deletions during editing, constraining the robustness of lineage tree reconstruction^17^.

Here, we introduce eTRACER, a novel CRISPR-Cas9 lineage tracer that targets endogenous 3’UTRs as evolving barcodes. This design facilitates efficient editing and robust barcode recovery via poly(A)-based single-cell RNA-seq (scRNA-seq) or high-resolution spatial transcriptomics (e.g., Stereo-seq^34^). Notably, eTRACER achieves exceptional spatiotemporal lineage resolution. Importantly, we have comprehensively demonstrated that the editing of neutral 3’UTRs has minimal impact on both the expression of endogenous target genes and cell fitness. We apply eTRACER to record the spatiotemporal lineages of lung cancer cells (*EGFR^L858R^;Trp53^-/-^*) under CD8^+^ T cell-mediated immunotherapy. The integration of longitudinal and multimodal lineage tracing enables a systematic dissection of evolutionary adaptation to immunotherapy and provides a rigorous test of the plasticity-first hypothesis of tumor evolution.

## Results

### Design of a multimodal lineage tracer eTRACER

The eTRACER system comprised 7 constitutively-expressed sgRNAs (cgRNAs) and 8 doxycycline (Dox)-inducible sgRNAs (igRNAs) specifically targeting the defined regions of 3’UTRs from 15 chosen endogenous genes (**Figure 1A, Figure S1A**). Upon Cas9 expression, these target sites can be edited sequentially, thus generating evolving barcodes for recording the division history of individual cells. Through scRNA-seq and targeted sequencing of the transcribed target sites, we can simultaneously profile transcriptomes and reconstruct phylogenetic trees at single-cell level (**Figure 1A**). To enable parallel tracking of clonal lineages, we engineered dual static barcode libraries (SB1 and SB2) into eTRACER by fusing a 14-bp randomized sequence to GFP and tdTomato, respectively (**Figure 1A, Figure S1B**).

**Figure 1.**
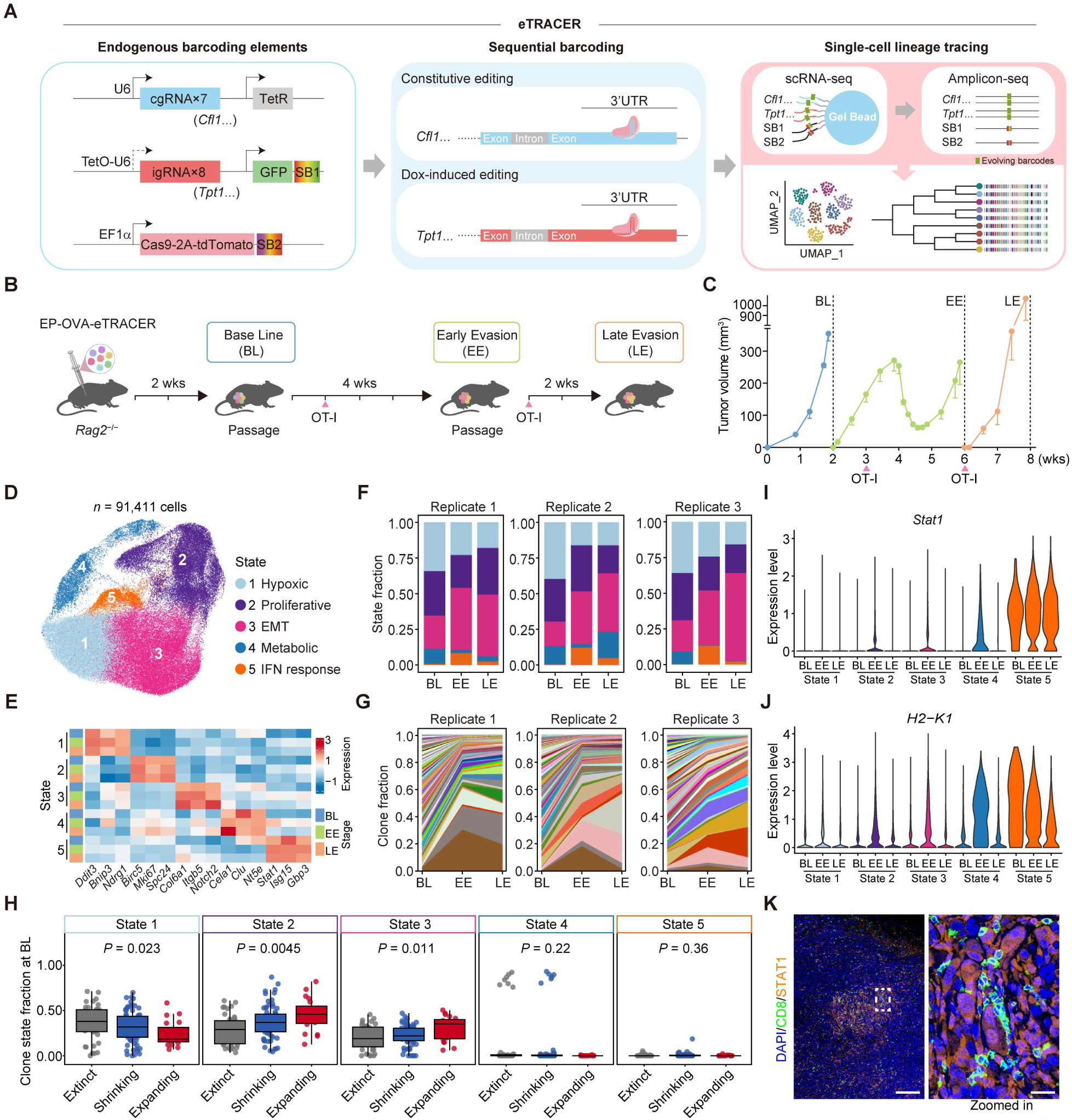
eTRACER enables longitudinal tracking of tumor states and clonal lineages during CD8⁺ T cell-mediated immunotherapy. (A) Schematic illustration of the eTRACER lineage tracing system. (B) Longitudinal transplantation of EP-OVA-eTRACER tumors over OT-I CD8⁺ T treatment. 5 × 10^4^ cells were implanted into *Rag2^-/-^* mice (Base-Line, BL, 2 weeks). Tumors were then serially passaged into two subsequent stages: Early-Evasion (EE, 4 weeks, OT-I at week 1) and Late-Evasion (LE, 2 weeks, OT-I at day 1). Doxycycline (1 mg/mL) was given continuously starting from week 1 of BL. scRNA-seq, spatial transcriptomics, and barcode sequencing were performed on samples from all three stages. (C) Growth curves of EP-OVA-eTRACER tumors over three stages (BL, EE and LE). Mean±s.e.m. (D) UMAP visualization of scRNA-seq data generated from tumors at the BL (n = 3), EE (n = 6) and LE (n = 6). (E) Heatmap showing marker gene expression for tumor cell states at individual stages. (F) Stacked bar charts showing the cell state fractions at each stage across three replicates. (G) Muller plots showing the clone fractions of top 173 clones at each stage across three replicates. (H) Association between the proportion of each cell state in the BL clones (≥ 20 cells) and clonal fate groups, assessed by the Kruskal-Wallis test. (I-J) Expression levels of *Stat1* (I) and *H2-K1* (J) across different stages and states. (K) Representative immunofluorescence images showing STAT1 and CD8 expression in EP-OVA-shCtrl tumor samples at day 21 post-OT-I injection (see Figure S19A). Localized view (DAPI/CD8/STAT1) of the day 21 sample image (DAPI/KRT8/CD8/STAT1) from Figure S8. Scale bars represent 200 µm and 20 µm (zoomed-in view), respectively.

To assess the potential impact of eTRACER on host cells, we introduced eTRACER into *EGFR^L858R^;Trp53^-/-^* (EP) mouse lung cancer cells. When cultured with Dox, these EP-eTRACER cells did not show notable changes in target gene expression or cell viability (**Figure S1C-D**). We further confirmed this in a panel of mouse cell lines including NIH/3T3, non-small cell lung cancer (*Kras^G12D^;Lkb1^-/-^*, KL), small cell lung cancer (*Rb1^-/-^;Trp53^-/-^*, RP), hepatocellular carcinoma (Hepa1-6), breast cancer (4T1) and melanoma (B16) cells (**Figure S1E-P**). Notably, eTRACER-induced deletions averaged approximately 4 bp, markedly shorter than the 8-10 bp deletions generated by current state-of-the-art CRISPR/Cas9 lineage tracers using exogenous tandem targets^21,23^ (**Figure S2A-C**). As expected, the 15 sgRNAs we designed exhibited gradient editing efficiencies in EP cells, among which the 8 igRNAs showed minimal leakiness without Dox induction (**Figure S2D**). Strong correlation between the number of unique evolving barcodes and the number of all edited reads indicated the high editing diversity of eTRACER (**Figure S2E**). These editing patterns were further confirmed in NIH/3T3 cells (**Figure S2F-I**). Together, these basic characteristics of eTRACER demonstrate its minimal impact on host cells and broad potential for probing cell state and lineage evolution across biological contexts.

### Tumor clonal and state dynamics during CD8^+^ T cell-mediated cytotoxicity

To investigate the adaptive evolution of tumor cells under immune killing, we used the OVA/OT-I system, a well-established, effective and clean model of CD8^+^ T cell-mediated cytotoxicity^35–37^. In this system, cancer cells with antigen ovalbumin (OVA) expression on the cell surface can be specifically killed by OT-I CD8^+^ T cells (OT-I cells hereafter) through recognizing the MHC-I/OVA complex. We engineered EP-eTRACER cells to stably express OVA (EP-OVA-eTRACER) and confirmed that the response of EP-OVA-eTRACER cells to OT-I cell killing was not affected by eTRACER *per se* (**Figure S3A-B**). We then performed longitudinal subcutaneous transplantation followed by two adoptive transfers of OT-I cells into *Rag2*^-/-^ mice (**Figure 1B**). Following the first injection of OT-I cells, tumors regressed dramatically but recurred approximately 3 weeks post-treatment. Moreover, re-injection of OT-I cells failed to inhibit tumor growth (**Figure 1C**), indicating that these serially-transplanted tumor cells had escaped OT-I cell-mediated cytotoxicity. We collected samples at the Base Line (BL, before OT-I cell injection), Early Evasion (EE, 3 weeks after first OT-I cell injection) and Late Evasion (LE, 2 weeks after second OT-I cell injection) stages for scRNA-seq, spatial transcriptomic profiling, and static/evolving barcode sequencing (**Figure 1A,C and Supplementary Table 1**).

Following standard quality control, we obtained 91,411 cells with high-quality scRNA-seq data (**Figure 1D**). Single-cell expression of the 15 endogenous target genes was consistent with *in vitro* bulk RNA-seq data (**Figure S3C-D**). Unsupervised clustering identified five heterogeneous tumor cell states^38,39^: Hypoxic (state 1), Proliferative (state 2), EMT (state 3), Metabolic (state 4), and Interferon response (state 5) (**Figure 1D-E, Figure S3E-F**). Notably, state 3 was enriched following OT-I cell treatment and remained predominant upon re-injection, whereas states 1, 2, and 4 were suppressed. State 5 showed a transient increase at EE but declined at LE, indicating dynamic trajectories during immunotherapy (**Figure 1F**).

Clone analysis based on static barcodes (**Figure S4A-C**) showed that only 10.5% of tracked clones (293 of 2,796) present at BL survived through the EE or LE stages, with the vast majority becoming extinct or falling below detection after therapy (**Figure 1G and Methods**). This pattern was consistent with the marked tumor shrinkage following the first dose of OT-I therapy (**Figure 1C**). The clonal fractions displayed only a weak correlation between BL and EE (**Figure S4D**), but a strong correlation between EE and LE (**Figure S4E**), suggesting establishment of immune escape and subsequent relaxation of selective pressure during the second dose of OT-I therapy.

To further probe the relationship between state compositions and fitness at clonal level, the top 173 clones (each > 15 cells) were stratified as three fate subgroups (expanding, shrinking, or extinct) based on the changes in clonal frequency during immunotherapy (**Figure S5 and S6**). Notably, all expanding clones had multi-state compositions including states 1, 2, 3, and 5 cells (**Figure S6**). In contrast, single-state clones, dominated by state 4 cells, were shrinking or became extinct during immunotherapy (**Figure S6**). These findings imply that cellular state plasticity might facilitate adaptation to immunotherapy. At clone level, the proportions of state 1 at BL were negatively associated, whereas the proportions of states 2 and 3 at BL were positively associated with the clonal fate from BL to EE (**Figure 1H**). These results suggest that the pre-existing clonal state heterogeneity dictates the clonal fate under immune stress, in line with an *in vitro* study revealing diverse cell fates emerging upon drug treatment in genetically identical cells^40^. Interestingly, while shrinking and expanding clones exhibited distinct state compositions at BL, these differences vanished at EE and LE stages (**Figure S7A-B**). Moreover, state compositions remained remarkably stable throughout EE/LE stages, with state 3 cells consistently representing the dominant population (**Figure S7A-B**). These data indicate that by the EE/LE stage, the surviving clones converge into stationary states adapted to immune pressure. Strikingly, the surviving clones across the three biological replicates showed high stochasticity, with only 27% recurring in at least two replicates (**Figure 1G**). Single-cell copy number variation (CNV) analysis using inferCNV^41^ confirmed that the overall CNV profiles were similar across clones, although certain cell states such as state 4 exhibited distinct CNV patterns (**Figure S7C**). Together, these findings suggest that evasion of OT-I cell-mediated cytotoxicity arises not from pre-existing clonal genetic heterogeneity, but from pre-existing cellular state plasticity.

We noted that state 5, characterized by activation of the JAK-STAT1 pathway and antigen presentation genes (**Figure 1I-J, Figure S3E-F and S7D**), increased markedly from BL to EE during immunotherapy, but subsequently declined from EE to LE (**Figure 1F**), consistent with an acute immune response. Immunofluorescence staining confirmed substantial co-localization of OT-I cells and state 5 tumor cells, supporting direct cytotoxic targeting by OT-I cells (**Figure 1K, Figure S8**). State 4 also decreased significantly following immunotherapy (except in replicate 2, where they rebounded at LE) (**Figure 1F**) and showed transient upregulation of the JAK-STAT1 pathway and antigen-presentation genes at EE (**Figure 1I-J, Figure S7D**).

### Phylogeny-resolved cell-state transitions during CD8⁺ T cell cytotoxicity

eTRACER-enabled simultaneous single-cell phylogenetic reconstruction and transcriptomic profiling allows systematic mapping of tumor state transitions driving immune evasion (**Figure 2, Figure S9 and S10**). The gradient editing efficiencies across different targets (**Figure 2A**), high recovery rates of target sites by scRNA-seq (median = 94%, **Figure S9A**), extensive editing events per cell (mean = 15, **Figure S9B-E**) and high diversity of evolving barcodes (> 80% of individual cells uniquely marked, **Figure S9F**) collectively enable the reconstruction of high-fidelity and high-resolution single-cell phylogenies (**Figure S10**). As expected, cells within the same clone (defined by static barcodes) exhibited smaller editing distances compared to those from distinct clones (**Figure 2B, Figure S9G**). Using an integrated phylogenetic tree reconstructed by maximum likelihood^42^ from the 30 most abundant clones (100 randomly sampled cells per clone), 92% of cells can be correctly separated in phylogeny by their clonal origins (**Figure 2C**). Bootstrapping analysis demonstrated high robustness of phylogenetic reconstruction, with the median confidence score ranging from 79% to 90% across the top 10 clones (**Figure S9H**). Compared with previous state-of-the-art CRISPR-Cas9 lineage tracers^21,23^, eTRACER phylogenies exhibited a 2- to 3-fold increase in the number of internal branching events (**Figure 2D, Figure S9I**), demonstrating its superior phylogenetic resolution.

**Figure 2.**
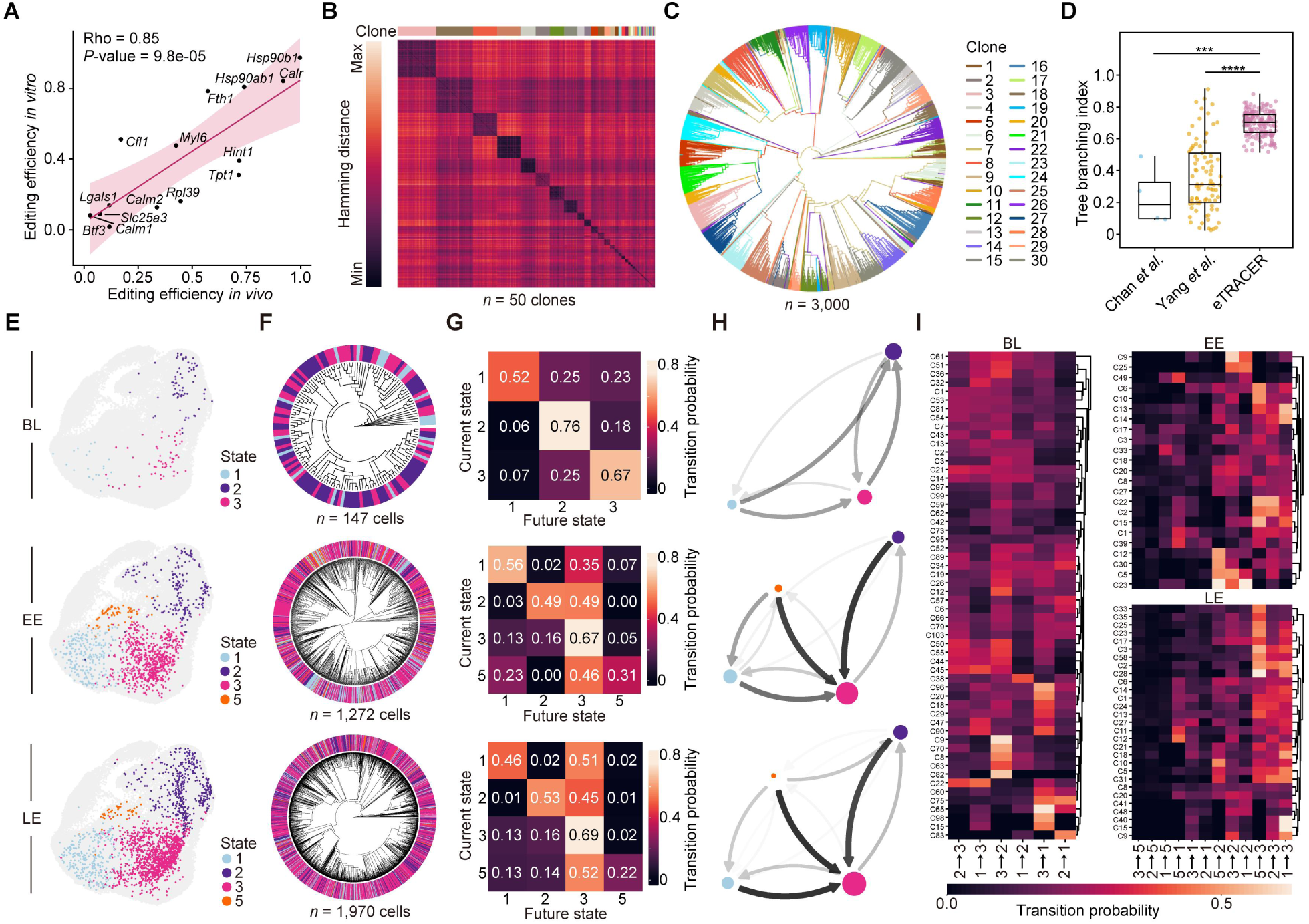
Quantifying cell-state transitions using single-cell phylogenies. (A) Correlation between *in vivo* (single cell) and *in vitro* (bulk) editing efficiencies at target sites. Spearman’s rho and *P-*value are shown. (B) Heatmap showing distance of evolving barcodes within clones at the EE stage. Representative data from replicate 2. (C) A single-cell phylogeny reconstructed using 3,000 randomly sampled cells from the top 30 clones (100 random cells for each clone). Clone identities of individual cells are visualized. (D) The branching index of eTRACER lineage trees (n = 173 clones, each >15 cells) and CRISPR-Cas9 lineage trees from previous studies (Chan *et al.*^23^, n = 4 clones; Yang *et al.*^21^, n = 85 clones). Wilcoxon test. (E-G) UMAP visualization (E), single-cell phylogenies (F) and state-transition probabilities between any two states (G) of clone 2 across the BL, EE, and LE stages. (H) State-transition networks in clone 2 across the BL, EE, and LE stages. Arrows indicate the direction of transition, and the thickness/color intensity of arrows indicate the transition probabilities. Circle size represents proportion of each state in the clone. (I) Heatmap showing state transition probability within individual multi-state clones at the BL, EE, and LE stages. Clones with ≥ 100 cells are shown.

With single-cell phylogenies alongside terminal cell states annotated by scRNA-seq data (**Figure 2E-F, Figure S10**), we next sought to construct a quantitative map of cell-state transitions. We observed extensive phylogenetic intermixing of cell states in 90% of clones (155 out of the top 173 clones) (**Figure S10**), indicating low state heritability and profound transcriptomic plasticity (**Figure S11**). We next used PATH^43^ to quantify the transition probabilities between any two cell states in each clone (**Figure 2G-I**). Taking clone 2 as an example, it harbored multi-state compositions including states 1, 2, and 3 cells at BL and states 1, 2, 3, and 5 cells at EE and LE (**Figure 2E**). Prior to therapy (BL), the state transitions within clone 2 were mainly state 1-to-2, 1-to-3 and 3-to-2, while the probabilities of reversal transitions (state 2-to-1, 3-to-1 and 2-to-3) were also high (**Figure 2H**), implying highly dynamic transitions and strong plasticity. However, after therapy (EE and LE), states 1, 2, and 5 exhibited pronounced directional transitions toward state 3 (**Figure 2H**), consistent with the overall increase of state 3 after immunotherapy (**Figure 1F**). This indicates that non-dominant phenotypes leverage phenotypic plasticity to convert into a dominant state (state 3), thereby bolstering their adaptation to immune pressure. State 3 exhibited the highest self-transition probability at both EE and LE stages (**Figure 2G**), reflecting enhanced state heritability following treatment. This finding is consistent with the stabilization of state 3 from EE to LE stages (**Figure 1F, Figure S7A-B**). Such state-transition pattern was indeed conserved across multi-state clones at three stages (**Figure 2I**). Therefore, our high-resolution phylogenies and coupled longitudinal scRNA-seq data generated by eTRACER demonstrate that the state 3 (EMT) serves as the crucial cell fate for immune escape.

### Spatially-resolved lineage tracing reveals tumor architectural adaptation

We next performed *in situ* lineage barcode capture via Stereo-seq^34^ using tumor samples having matched scRNA-seq data across all three stages (**Figure 3A**). Three samples from each stage were co-arrayed on a single Stereo-seq chip (**Figure 3A**). Through amplicon-seq of the static and evolving barcodes with Stereo-seq cDNA libraries, we were able to simultaneously resolve spatial transcriptomic profiles and lineage relationships at near-single-cell resolution (25 μm, namely bin 50, **Figure 3A, Figure S12A-B**). Spatial cell types were annotated using the scRNA-seq data as reference, encompassing tumor, immune and stromal cells (**Figure 3B**). In line with scRNA-seq data, tumor cells at BL mainly consisted of states 1, 2, and 3, whereas those at EE/LE were dominated by states 3 and 2 (**Figure 3C, Figure S12C**). Spatial neighborhood analysis revealed closer proximity between states 1 and 3, as well as between states 2 and 3 (**Figure 3D**). Notably, a pronounced spatial stratification of states 1, 2, and 3 was observed, where the Hypoxic state (state 1) primarily localized at the tumor interior, the Proliferative state (state 2) at the tumor margin and the EMT state (state 3) at an intermediate zone (**Figure 3E, Figure S12D**). This stratified spatial pattern was further validated by immunofluorescence analysis (**Figure S12E-F**). Spatial clone mapping using static barcodes also revealed that cells from the same clone were more aggregated, suggesting localized clonal expansions in general (**Figure 3F-G, Figure S13A**). We observed a strong association between within-clone state composition and the clone distance to tumor margin, where the proportion of state 2 increased and the proportion of state 3 decreased as clones shifted toward the tumor margin (**Figure S13B-C**). These findings indicate a strong influence of local microenvironment on tumor clonal states^38^.

**Figure 3.**
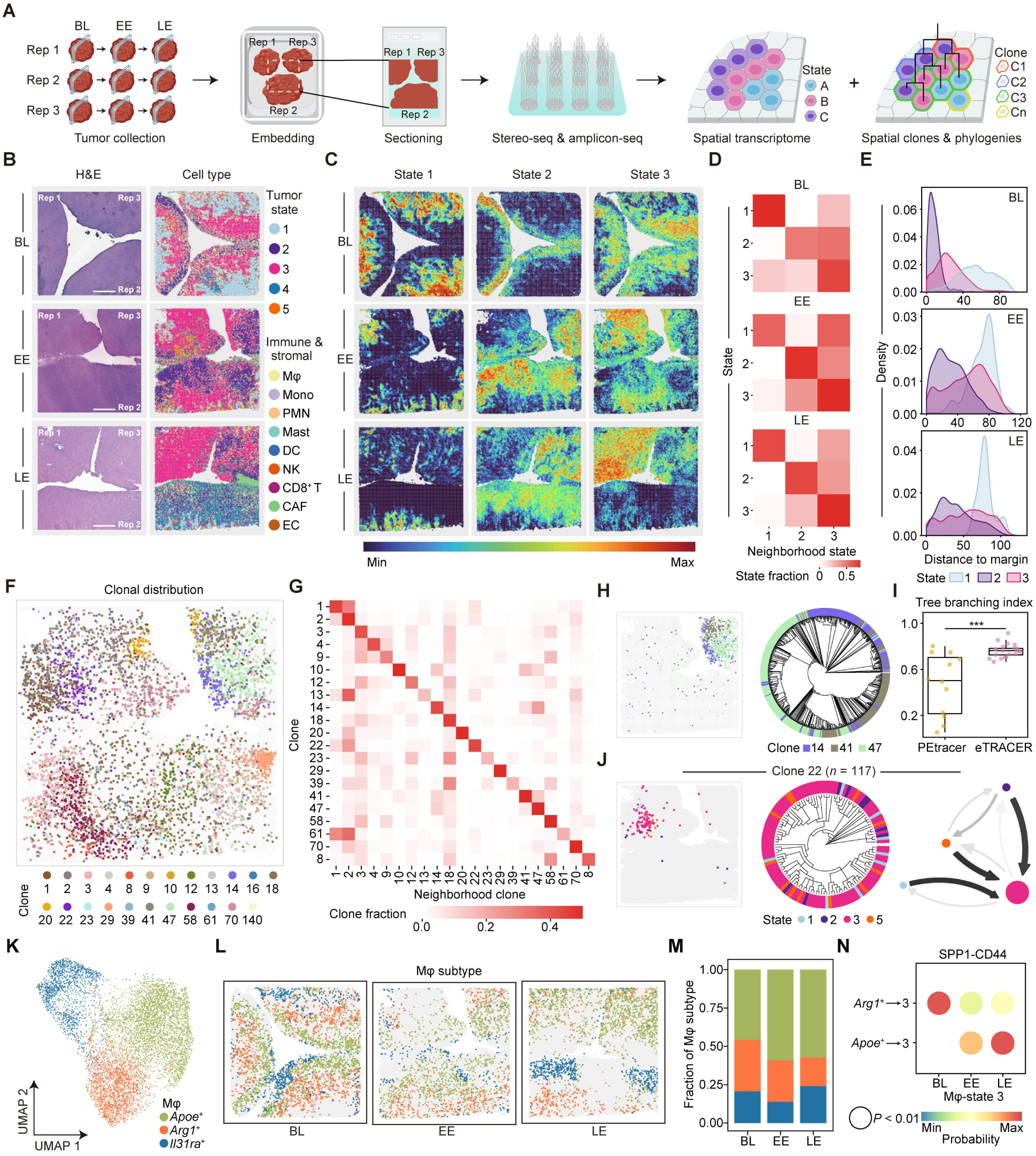
Spatially-resolved lineage tracing reveals architectural organization and directional transitions of tumor cell states. (A) Schematic illustration of spatial transcriptomic and lineage barcode profiling. Rep, replicate. (B) Representative H&E images and spatial mapping of tumor cells, immune and stromal cells at the BL, EE and LE stages. Mφ, macrophage; Mono, monocyte; PMN, polymorphonuclear neutrophil; DC, dendritic cell; NK, natural killer cell; CAF, cancer-associated fibroblast; EC, endothelium cell. Scale bar, 1 mm. (C) Spatial mapping of state 1 (Hypoxic), state 2 (Proliferative) and state 3 (EMT) at the BL, EE and LE stages. (D) Heatmap showing the spatial neighborhood compositions and fractions of indicated states. (E) Quantitative analyses of spatial distances to the tumor margin for states 1, 2, and 3. The unit of X-axis is bin 50 (25 μm). The area under the curve for each state sums up to 1. (F) Spatial clone mapping at the EE stage. Clones with > 20 cells were included and those with > 500 cells were downsampled to 500. (G) Neighborhood analysis of tumor cell clones at the EE stage. (H) Spatial mapping of three representative tumor cell clones. A spatial phylogeny reconstructed using 625 bin 50s from clone 14, 41, and 47 based on Stereo-seq data analyses. (I) The branching index of eTRACER spatial lineage trees (n = 21 clones) was compared with that of PEtracer spatial lineage trees (n = 12 clones)^31^. In both datasets, phylogenetic trees were constructed using the maximum likelihood method. Clones containing more than 2,000 cells were downsampled to 2,000 cells. Wilcoxon test. (J) Spatial locations, phylogenetic trees and state-transition networks of clone 22. (K) UMAP visualization of macrophage subtypes. (L) Spatial mapping of macrophage subtypes at the BL, EE and LE stages. (M) The fractions of macrophage subtypes at the BL, EE and LE stages. (N) Quantitative probabilities of SPP1-CD44-mediated macrophage-state 3 cell interactions through cell-cell communication analyses. *P*-value < 0.01.

To further quantify spatial state transitions, we reconstructed spatially-resolved phylogenies using the evolving barcodes captured by Stereo-seq (**Figure S14A-G**). Integrative phylogenetic reconstruction by merging multiple clones confirmed the high accuracy of the lineage trees (**Figure 3H**). Cells within the same subclone exhibited prominent spatial clustering, consistent with their high phylogenetic relatedness (**Figure S14H**). Notably, as compared to PEtracer^31^, a primer editing and imaging readout-based spatial lineage tracer, eTRACER-reconstructed spatial lineage trees exhibited higher phylogenetic branching (**Figure 3I**) and longer tree depth (**Figure S14F and S15**). Importantly, PATH analysis with the spatial lineage trees also revealed pronounced state transitions from state 1 to 3 and state 2 to 3 (**Figure 3J, Figure S14I and S15**), as in our phylogeny-resolved scRNA-seq data (**Figure 2H-I**).

Spatial transcriptomics also showed that macrophages (Mφ) constituted the predominant immune cell population within the tumor microenvironment (TME) both before and after therapy (**Figure S16A-B**), with evident spatial co-localization with the state 3 cells (**Figure S16C-F**). Macrophages can be further classified into three subtypes based on their transcriptomic profiles: *Apoe^+^*, *Arg1^+^*, and *IL31ra^+^,* respectively^44^ (**Figure 3K-M, Figure S16G-H**). Through spatial analysis of cell-cell communications (CCC), we identified significant SPP1-CD44 ligand-receptor interactions between macrophages and state 3 cells (**Figure 3N**). Interestingly, there was a dramatic shift from *Arg1^+^* macrophages at BL to *Apoe^+^* macrophages at EE/LE interacting with CD44^+^ state 3 cells (**Figure 3N**), in line with the expansion of *Apoe^+^* macrophages after CD8^+^ T cell treatment (**Figure 3M**). These data indicate that CD8^+^ T cell-mediated immunotherapy also facilitates the reprogramming of macrophages. The CCC analysis also revealed interactions mediated by state 3 tumor cells, potentially driving the recruitment and polarization of macrophages via the MIF-CD74/CD44 and CSF1-CSF1R axes^45,46^ (**Figure S16I**). These findings underscore the co-evolutionary dynamics between tumor cells and macrophages during CD8^+^ T cell-mediated immunotherapy.

### AP-1 mediates the phenotypic adaptation and stabilization

To identify the key regulator underlying the adaptation of state 3 cells during immunotherapy, we performed single-cell multiome sequencing (scMultiome)^47^ to jointly profile the transcriptome, chromatin accessibility and lineage barcodes (**Figure 4A-B**). scMultiome identified the same five tumor cell states as those in the scRNA-seq data (**Figure 1D**, **Figure 4C, Figure S17A-C**). Simultaneous recovery of the lineage barcodes further demonstrated the high fidelity of inferred lineage relationships (**Figure 4D, Figure S17D**). Consistently, states 1 and 2 showed clear transitions to state 3 according to phylogenetic tree from scMultiome data (**Figure S17E-G**). Transcription factor (TF) regulon analysis revealed that *Jun* and *Fosb*, the core components of the activator protein-1 (AP-1), were specifically enriched within state 3 (**Figure 4E**). This was further confirmed by the scRNA-seq data (**Figure 4F, Figure S17H-I**). As expected, EMT-associated pathways were highly enriched among the AP-1 target genes (hereafter referred to as the AP-1 signature) in state 3 (**Figure 4G-H**). Consistently, both *Jun/Fosb* and EMT-associated pathways were significantly upregulated in expanding clones as compared to extinct clones in scRNA-seq data (**Figure S17J-K**). The AP-1 signature score was significantly elevated at EE and remained stable throughout LE, particularly in state 3 (**Figure 4I, Figure S17L**). Compared with BL, tumor cells from both EE and LE exhibited specific enrichment of the AP-1 binding motif (**Figure 17M**) and increased chromatin accessibility at AP-1 signature loci (**Figure S17N-O**). These data together uncover that AP-1 is the key regulator of state 3, sustaining the AP-1 signature via transcriptional memory following longitudinal immunotherapy.

**Figure 4.**
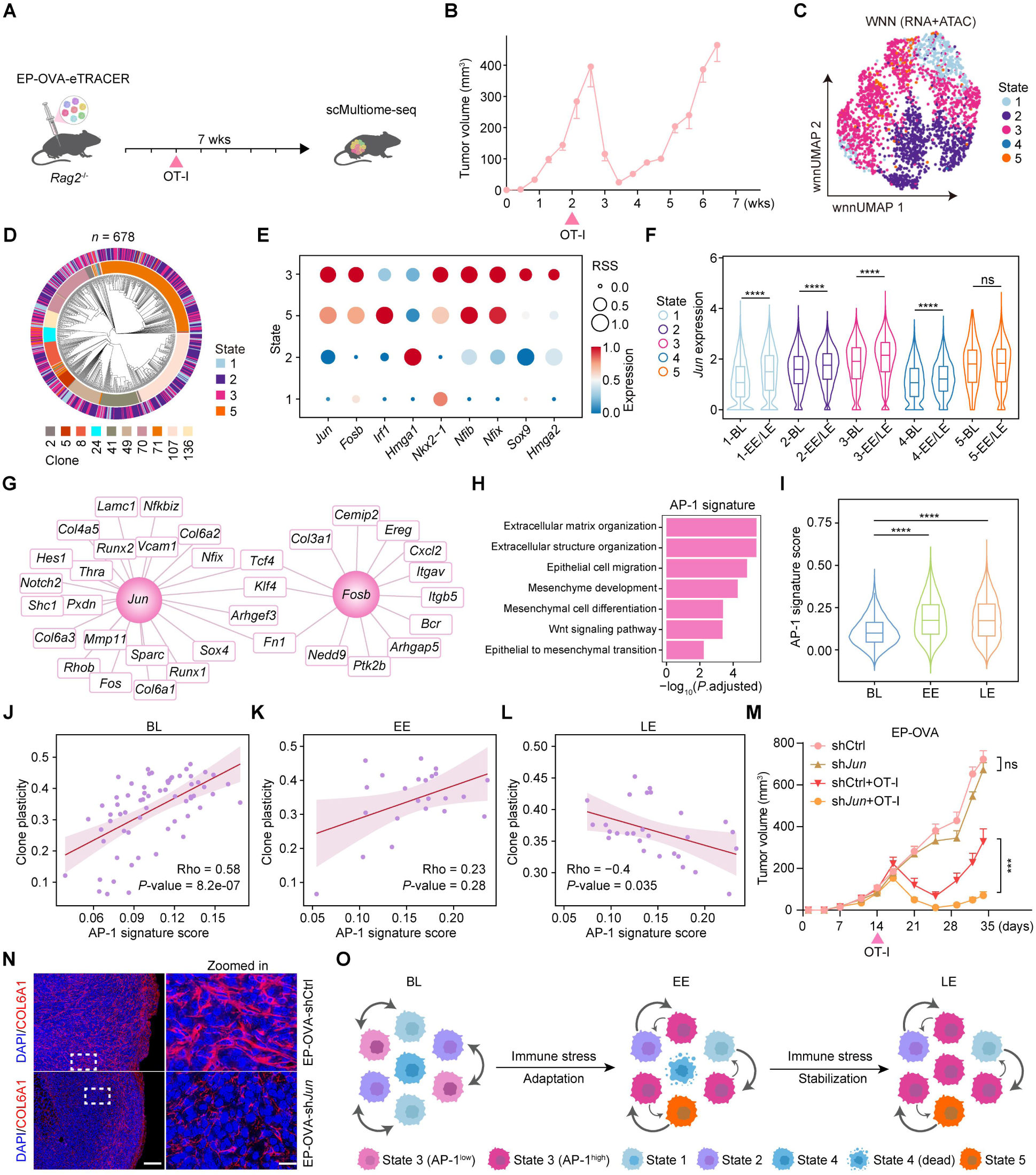
Single-cell multiomic analysis identifies AP-1 as the key regulator of phenotypic adaptation and stabilization. (A) Schematic illustration of sample collection for scMultiome-seq. 50,000 EP-OVA-eTRACER cells were subcutaneously transplanted into *Rag2^-/-^* mice. Dox (1 mg/mL) administered in drinking water at 1 week post-engraftment. At 2 weeks post-engraftment, 1 × 10^6^ OT-I cells were injected intravenously and tumors were harvested for scMultiome-seq at 7 weeks post-engraftment. (B) Tumor growth curve of EP-OVA-eTRACER. Mean±s.e.m. (C) UMAP visualization of integrative scMultiome data (n = 2,568 cells) based on weighted nearest-neighbor (WNN) analysis. (D) Single-cell phylogeny reconstructed based on evolving barcodes from 678 cells belonging to clones with > 15 cells, as identified by scMultiome-seq. In the phylogenetic tree, the outer and inner rings represent cell state and clone identity, respectively. (E) Dot heatmap showing the expression and regulon specificity scores (RSS) of transcription factors in cell states identified through scMultiome analyses. (F) Statistical analysis of the expression of *Jun* in scRNA-seq data of different tumor cell states at the BL and EE/LE stages. Wilcoxon test. (G) The representative target genes of AP-1 (*Jun*/*Fosb*). (H) Pathway enrichment analysis of AP-1 (*Jun*/*Fosb*) target genes using scMultiome data. (I) AP-1 signature scores in single-cell level across the three stages. Wilcoxon test. (J-L) Correlation between clonal plasticity and AP-1 signature score at the BL (J), EE (K), and LE (L) stages. Spearman’s rho and *P-*value are shown, \**P* < 0.05; \*\**P* < 0.01; \*\*\**P* < 0.001; \*\*\*\**P* < 0.0001; ns, not significant, *P* ≥ 0.05. (M) Tumor growth curves of EP-OVA-shCtrl or EP-OVA-sh*Jun* cells in *Rag2^-/-^* mice (n ≥ 3 mice per group). 5 × 10^4^ EP-OVA-shVector or EP-OVA-sh*Jun* cells were subcutaneously inoculated into *Rag2*^-/-^ mice. Two weeks post-engraftment, the mice received an intravenous injection of 1 × 10^6^ OT-I cells. Tumor volumes were monitored for a total of 5 weeks. Two-sided Student’s t-test, \**P* < 0.05; \*\**P* < 0.01; \*\*\**P* < 0.001; \*\*\*\**P* < 0.0001; ns, not significant, *P* ≥ 0.05. Mean±s.e.m. (N) Representative immunofluorescence images showing COL6A1 expression in tumor samples at day 3 post-OT-I injection (see Figure S19A). DAPI and COL6A1 channels extracted from the triple-stained images (DAPI/KRT8/COL6A1) in Figure S19D. Scale bars represent 200 µm and 20 µm (zoomed-in view), respectively. (O) A plasticity-first model of tumor immune evasion indicates lineage dynamics of phenotypic adaptation and stabilization.

We next asked whether AP-1 plays a key role in the state transitions and the stabilization of state 3. The high-resolution single-cell phylogenies with terminal cell state annotated allow us to quantify the cell-state plasticity at clone level (**Figure S10**), using Fitch-Hartigan maximum-parsimony algorithm^48,49^. We observed a strong positive correlation (Spearman’s rho = 0.58, *P*-value = 8.2 × 10^-7^) between AP-1 signature score and within-clone state plasticity prior to therapy at BL (**Figure 4J**). While this correlation weakened at EE (**Figure 4K**), a significant negative correlation emerged at LE (Spearman’s rho = -0.4, *P*-value = 0.035, **Figure 4L**). These findings suggest that low level AP-1 likely unlocks cellular plasticity, while sustained upregulation of AP-1 during immunotherapy stabilizes the adaptive phenotype (state 3) and constrains overall plasticity. Indeed, we found higher AP-1 signature was associated with higher plasticity of states 1, 2, and 5 but lower plasticity of state 3 (**Figure S17P**). Given the convergent transitions toward state 3 observed at the EE/LE stages (**Figure 2H-I**), our results indicate that AP-1 orchestrates this convergence while stabilizing the state 3 phenotype.

Interestingly, spatially diffused clones exhibited elevated expression of AP-1 (*Jun* and *Fosb*) as compared to spatially localized clones (**Figure S18**). These data suggest that AP-1 might promote tumor cell migration, in line with the enrichment of the AP-1 target genes in epithelial cell migration and EMT pathways (**Figure 4H**). To further validate the critical role of AP-1 in immune evasion, we generated *Jun*-knockdown tumor cells and assessed their response to OT-I T cell therapy *in vivo* (**Figure S19A-C**). Notably, *Jun* depletion significantly restored tumor sensitivity to T cell-mediated killing, leading to enhanced tumor control (**Figure 4M**). Indeed, *Jun* depletion significantly reduced state 3 cells according to marker-specific immunofluorescence staining (**Figure 4N, Figure S19D**).

Collectively, these findings establish AP-1 as a key mediator of stable immune evasion during immunotherapy, revealing a transcriptional mechanism of phenotypic adaptation within the plasticity-first framework (**Figure 4O**).

## Discussion

In this study, we rigorously test and mechanistically investigate the plasticity-first hypothesis of tumor evolution using eTRACER, a high-resolution, spatiotemporally resolved multimodal lineage-tracing system. In a CD8⁺ T cell-mediated immunotherapy model of EGFR-mutant lung cancer, we show that tumor phenotypic plasticity precedes stable immune evasion, while AP-1 upregulation upon immunotherapy consolidates adaptive state transitions and drives long-term resistance to persistent immunotherapy.

Although immune checkpoint blockade represents a groundbreaking advance in cancer treatment, EGFR-mutant lung adenocarcinoma patients show markedly poor responsiveness to anti-PD-1/PD-L1 immunotherapies^50,51^. By engineering EGFR-mutant lung cancer cells (*EGFR^L858R^;Trp53^-/-^*) to specifically express the antigen OVA and treating them with OT-I CD8^+^ T cells *in vivo*, we are able to dissect the intrinsic mechanisms of immune evasion in EGFR-mutant lung cancer. Using eTRACER to delineate the tumor spatiotemporal lineages and state evolution during immunotherapy, we first observe highly stochastic clone fates, with only 27% of surviving clones recurring in at least two biological experimental replicates. However, we find that the proportion of EMT cells within individual clones correlates with clonal survival and expansion. These observations argue against the conventional clonal selection model^1–3^ where immune evasion is genetically predetermined and selected from pre-existing clonal heterogeneity. Instead, our systematic and quantitative analyses with phylogeny-resolved scRNA-seq, spatial transcriptomics, and scMultiome datasets reveal high phenotypic plasticity prior to therapy, characterized by highly dynamic and reversible cell-state transitions. Surprisingly, we find that immunotherapy induces convergent state transitions to a dominant EMT state with antigen presentation defect, which is subsequently stabilized throughout the course of treatment.

We further identify AP-1 transcription factor as a key regulator driving the long-term tumor phenotypic adaptation in response to immunotherapy. First, we demonstrate that AP-1 is a central regulator of the EMT state. Crucially, the expression of AP-1-regulated genes and their chromatin accessibility increase significantly during initial immunotherapy and are maintained throughout longitudinal treatment. Our phylogeny-resolved single-cell multiomic data show that the heritability of the EMT state increases and persists during immunotherapy, reflecting a marked decline in phenotypic plasticity post therapy. These data suggest that AP-1 stabilizes the EMT state via transcriptional memory, in line with recent studies demonstrating that AP-1 mediates cellular memory of targeted therapy resistance^52,53^ and inflammation^54,55^.

The spatial architecture of tumor cell states within the tumor microenvironment may serve as a fundamental basis for tumor adaptation to therapeutic pressure^56^. By integrating spatial state and clonal mapping, we observe a distinct spatial stratification of tumor states with the EMT state occupying an intermediate niche, exhibiting significant co-localization and robust interactions with macrophages. Previous studies have reported that the expression and activation of AP-1 are regulated by multiple signals, ranging from intracellular oncogenic activation to extracellular cytokines and growth factors triggered by stress or inflammation^57^. We hypothesize that macrophages promote AP-1 activation in state 3 cells via SPP1-CD44 interactions, as supported by previous studies^58–60^. This finding underscores the capacity of tumor cells to exploit microenvironmental signals that drive their adaptive evolution.

Collectively, our comprehensive multimodal lineage tracing analyses demonstrate that the development of a stable immune-evasive phenotype arises from consolidated state transitions that leverage pre-existing cellular plasticity mediated by AP-1, supporting the plasticity-first hypothesis of immunotherapeutic resistance. We demonstrate that suppressing AP-1 can sensitize tumors to CD8^+^ T cell killing. Therefore, AP-1 inhibitor-based strategies to enhance adoptive cell therapy represent a promising avenue to overcome immunotherapy resistance, warranting future clinical investigations.

## Methods

### eTRACER design

eTRACER is comprised of three components: Cas9, sgRNAs (cgRNAs array and igRNAs array) and a dual static barcode system. The 15 sgRNAs for eTRACER were designed according to following criteria: 1) target genes are endogenous and highly expressed across various mouse organs^61^, to ensure efficient barcode detection; 2) cleavage sites are located in the 3’ UTR, 100-300 bp upstream of the 3’ end, to enable efficient amplicon-seq; 3) target sites should avoid conserved miRNA binding sites based on "miRBase" database^62^ to prevent disruption of endogenous gene expression; 4) target sequences are unique in the mouse genome when designed by the CRISPOR^63^; 5) the panel of 7 cgRNAs and 8 igRNAs should encompass high, medium, and low editing efficiencies to extend the tracing duration; and 6) sgRNA editing should have minimal impact on endogenous gene expression and cell viability. The cgRNAs were driven by U6 promoter whereas the igRNAs were driven by the Dox-inducible U6 promoter^64^, which contains the tet-operator (TetO) site allowing for the suppression of sgRNA transcription in the presence of tetracycline repressor (TetR). Both cgRNA and igRNA arrays were assembled using the Golden Gate Assembly Kit (NEB, #E1601S). A 14-bp static barcode library was cloned into the pCDH-EF1-Cas9-P2A-tdTomato and pCDH-TetO-U6-igRNAs-EF1-GFP plasmids as previously described^65^. The primers used for plasmid construction were listed in Supplementary Table 2.

### In vitro and in vivo eTRACER experiments

Mouse cell lines including NIH/3T3, EP (*EGFR^L858R^;Trp53^-/-^*, lung adenocarcinoma), KL (*Kras^G12D^*;*Lkb1^-/-^*, lung adenocarcinoma), RP (*Rb1^-/-^*;*Trp53^-/-^*, small cell lung cancer), Hepa1-6 (hepatocellular carcinoma), 4T1 (breast cancer) and B16 (melanoma) were transfected with U6-cgRNAs-EF1-TetR-P2A-Puro, TetO-U6-igRNAs-EF1-GFP-SB1 and EF1-Cas9-P2A-tdTomato-SB2 to integrate the eTRACER components. Endogenous gene expression and cell viability were analyzed after 2 weeks of eTRACER transfection in the presence of 0.5 µg/mL Dox (MCE, #HY-N0565).

Both parental EP cells and EP-eTRACER cells were stably transfected with lentivirus vector expressing the OVA-Luciferase transgene. *In vitro* cell culture and *in vivo* mouse model experiments were conducted to analyze the potential impact of eTRACER on tumor growth and OT-I CD8*^+^* T cell-mediated killing. In details, OT-I CD8*^+^* T cells were isolated from the spleens of OT-I mice using the mouse CD8^+^ T cell isolation kit (STEMCELL Technologies), and resuspended with cRPMI medium (RPMI-1640 containing 10% FBS, 2 mM L-Glutamine, 55 μM β-mercaptoethanol, 10 ng/mL IL-2, 5 ng/mL IL-7, 5 ng/mL IL-15, 1 μg/mL anti-CD28) and cultured with plate-coated anti-CD3 (5 μg/mL) for 3 days. Subsequently, they were co-cultured with EP-OVA-eTRACER cells in ratios ranging from 0:1 to 8:1 for 24 hours, and 150 µg/mL luciferin (Beyotime, #ST196) was then added to each well, and luminescence was measured with an Envision plate reader (PerkinElmer). To test the impact of eTRACER on tumor growth and CD8^+^ T cell-mediated killing, 5 × 10^4^ EP-OVA or EP-OVA-eTRACER cells were subcutaneously inoculated into *Rag2*^-/-^ mice for 7 weeks. Dox (1 mg/mL in drinking water) was given at 1 week post-engraftment and 1 × 10^6^ OT-I cells were intravenously injected at 2 week post-engraftment and the tumor volume was monitored twice a week. For *in vitro* editing efficiency analysis, EP-eTRACER and NIH/3T3-eTRACER cells were cultured with Dox for two weeks and genomic DNA was extracted for amplicon-seq (Primers listed in Supplementary Table 3).

A multi-stage experimental system was established to track EP tumor evolution during OT-I-mediated immune surveillance. The workflow is as follows: 1) Baseline tumors were generated by engrafting 5 × 10^4^ EP-OVA-eTRACER cells subcutaneously in *Rag2^-^*^/-^ mice (n = 3) and the tumors were harvested 2 weeks later. This was designated as the BL (base line) stage. 2) BL-derived tumors were digested into single cells, using one half for serial transplantation into new *Rag2^-^*^/-^ mice (n = 6) and the other half for FACS sorting followed by scRNA-seq profiling. 1 × 10^6^ OT-I cells were intravenously injected at 1 week post-engraftment and tumors were harvested at 4 weeks post-engraftment. This stage was designated as early evasion (EE). 3) EE-derived tumors were digested into single cells, and used one half for serial transplantation into new *Rag2^-^*^/-^ mice (n = 6) and the other half for FACS sorting followed by scRNA-seq profiling. 1 × 10^6^ OT-I cells were intravenously injected at 1 day post-engraftment and tumors were harvested at 2 weeks post-engraftment. This stage was designated as LE (late evasion). Dox treatment (1 mg/mL in drinking water) started at 1 week post-engraftment at BL stage and continued throughout EE/LE stages. The scRNA-seq were performed using GFP^+^/tdTomato^+^ cancer cells isolated via FACS sorting from BL, EE, and LE tumors (n = 15) whereas Stereo-seq were performed using representative BL, EE and LE tumors (n = 3 each).

For scMultiome mouse experiment, 5 × 10^4^ EP-OVA-eTRACER cells were subcutaneously inoculated into *Rag2*^-/-^ mice for 7 weeks and the GFP^+^/tdTomato^+^ cancer cells were isolated via FACS sorting and subjected to scMultiome-seq. Dox (1 mg/mL in drinking water) treatment started at 1 week post-engraftment and 1 × 10^6^ OT-I cells were intravenously injected at 2 week post-engraftment. Tumor volumes were monitored every 3 days.

To validate AP-1 function, we first cloned a short hairpin RNA (shRNA) specifically targeting mouse *Jun* into the pLKO.1-U6-puro vector. The target sequence is: 5’-GCTAACGCAGCAGTTGCA AAC-3’. EP-OVA cells were transduced with lentiviral particles expressing either the *Jun*-targeting shRNA (sh*Jun*) or a non-targeting control vector (shVector). Stable cell lines were established following puromycin selection. Knockdown efficiency and downstream gene were verified by quantitative real-time PCR (qRT-PCR) to assess mRNA expression levels of *Jun* and *Col6a1* in the EP-OVA-shVector and EP-OVA-sh*Jun* cell lines. 5 × 10^4^ EP-OVA-shVector or EP-OVA-sh*Jun* cells were subcutaneously inoculated into *Rag2*^-/-^ mice. Two weeks post-engraftment, the mice received an intravenous injection of 1 × 10^6^ OT-I cells and were monitored for a total of 5 weeks. Tumor tissues from both groups were harvested at days 3, 7, and 21 post-OT-I cell transfer for subsequent analysis. Briefly, formalin-fixed paraffin-embedded (FFPE) sections were stained using a tyramide signal amplification (TSA)-based multiplex immunofluorescence kit (#AFIHC034, Hunan Aifang Biological Technology, Changsha, China) according to the manufacturer’s instructions. The panel of primary antibodies included: KRT8, DDIT3, BIRC5, F4/80 and CD8 (Hunan Aifang Biotechnology); COL6A1 (Cell Signaling Technology, #52395T); CELA1 (Thermo Fisher Scientific, #PA5-122103); STAT1 (Cell Signaling Technology, #14995S).

All mouse experiments complied with relevant ethical regulations and were approved by the Institutional Animal Care and Use Committee of the Center for Excellence in Molecular Cell Science, Chinese Academy of Sciences (SIBCB-2101008).

### Bulk ATAC-seq, Bulk RNA-seq, scRNA-seq, scMultiome-seq, Stereo-seq and Amplicon-seq

Bulk ATAC-seq was performed by LC-Bio Technology Co., Ltd. (Hangzhou, China) following standard protocols. Bulk RNA-seq (for parental EP cells), scRNA-seq and scMultiome-seq were performed by Berry Genomics Co., Ltd (Beijing, China) following standard protocols. Single tumor cell suspensions from individual mice were multiplexed using the 3’ CellPlex Kit (10x Genomics, #1000261) following manufacturer’s protocols to enable sample discrimination during scRNA-seq. Stereo-seq were performed by Beijing Novogene Technology Co., Ltd. following BGI protocol (v1.2 kit) and sequenced on MGI DNBSEQ sequencer.

Static and evolving barcode amplicon libraries were generated through targeted amplification of cDNA derived from the single-cell and spatial omics experiments (scRNA-seq, scMultiome-seq, Stereo-seq) using custom primers specific to barcode regions and target sites (Supplementary Table 4 and Table 5), and sequenced on the NovaSeq platform (Illumina) to generate 150bp paired-end reads.

### Bioinformatic analyses of bulk ATAC-seq, bulk RNA-seq, scRNA-seq, scMultiome-seq and Stereo-seq data

For bulk ATAC-seq data analyses, Fastp (v0.23.4)^66^ was used to obtain high-quality reads. Bowtie2 (v 2.5.4)^67^ was used to align the reads to the GRCm38 reference genome. MACS2 (v2.2.9.1)^68^ used for peak calling. DiffBind (v3.12.0)^69^ was used for the count table of consensus peaks normalization and differentially accessible peaks analysis. HOMER (v5.0)^70^ was used for motif enrichment analysis of the differentially accessible peaks with corresponding genomic positions as background. The AP-1 signature chromatin accessibility was visualized with the scaled normalized peaks.

For bulk RNA-seq data analyses, Cutadapt (v4.4)^71^ was used to trim adapters and obtained high-quality reads. HISAT2 (v2.2.1)^72^ was used to align the reads to the GRCm38 reference genome. Gene expression levels were quantified using featureCounts (v2.0.6)^73^.

For scRNA-seq data analyses, reads were aligned to the reference genome (mm10) and gene expression was quantified using CellRanger (v7.1.0). Cells with at least 1,000 genes and less than 10% mitochondrial genes were retained as high-quality cells. Genes expressed in < 3 cells were filtered out. DoubletFinder (v2.0.3)^74^ were used to remove mixed cells. A total of 91,411 high-quality tumor cells from the three time points in 15 mice (BL: n = 3, EE: n = 6 and LE: n = 6) were merged for downstream analysis. Using Seurat (v5.0.1)^75^, the cell count matrix was normalized, and 2,000 highly variable genes were selected for principal component analysis (PCA). To remove batch effects between samples, Harmony (v1.2.0)^76^ was applied, which demonstrates superior performance in distinguishing biological and technical variations. Tumor cell states were identified using the FindNeighbors function with the top 40 dimensions based on Harmony-corrected space, followed by the FindClusters function with a resolution of 0.15 and the Louvain algorithm. For single-cell visualization, non-linear dimensional reduction was performed using uniform manifold approximation and projection (UMAP). To annotate the biological functions of tumor states, Gene Ontology (GO) enrichment analysis was performed using the clusterProfiler package (v4.10.0)^77^ on the top 100 feature genes for each state (Supplementary Table 6), and Gene Set Variation Analysis (GSVA) was performed using the GSVA package (v1.50.5)^78^ on the hallmark gene set (mh.all.v2023.2.Mm.symbols.gmt). InferCNV (v1.18.1)^41^ was used to infer single-cell copy number variations. A total of 1,500 AT2-like cells were downsampled from the Jarod *et al*. dataset^79^ as the control group. For each tumor state, 1,500 cells were randomly downsampled from three time points. The analysis was performed with default parameters, except that the minimum average read count per gene was set to 0.1.

For scMultiome analyses, reads from four replicate tumor samples were aligned to the reference genome (mm10). Gene expression and chromatin accessibility quantification was performed using Cell Ranger ARC (v2.0.2). Cells with at least 1,000 genes and less than 20% mitochondrial genes were retained as high-quality cells. Genes expressed in < 3 cells were filtered out. A total of 2,568 high-quality tumor cells were retained. Cell state annotation was performed using Seurat’s FindTransferAnchors and TransferData functions, with the top 50 principal components, using the scRNA-seq data as reference. For gene regulatory network inference, tumor cell states are regulated by specific TFs and their target genes. SCENIC (v0.12.1)^80^ was used for scRNA-seq data. Co-expression modules between TF and candidate target genes were inferred using grnboost2. Modules with significant enrichment of the regulator’s binding motif across the target genes were identified and regulons with only direct targets were created using RcisTarget. The activity of each regulon in each cell was scored using AUCell. The SCENIC+ (v1.0a2)^81^ pipeline was used to analyze scMultiome data, using default parameter settings for quality control. Enhancer candidates as sets of co-accessible regions across tumor cell states were identified from scMultiome data using pycisTopic. Enriched TF-binding motifs were identified using pycisTarget. Linking TFs to candidate enhancers and target genes by integrating region accessibilities, TFs and target gene expressions. AP-1 signature scores were calculated using the Seurat AddModuleScore function.

For Stereo-seq analyses, reads from tumor tissues at three stages were aligned to the reference genome (mm10) and quantified for gene expression using the SAW pipeline^82^, which is tailored for preprocessing Stereo-seq data. Based on the adjacent staining tissue images, DNA nanoballs (DNBs) lacking associated tissue were excluded. To reduce noise from spatial sparsity and RNA diffusion, the raw count matrix was binned into a bin 50 matrix (with a diameter of 25 µm per bin 50) by aggregating gene expression from 50 × 50 DNBs. Bin 50s with less than 200 genes and genes expressed in fewer than 3 bin 50s were filtered out. 32,023 high-quality bin 50s were obtained at BL, 35,334 at the EE and 35,835 at LE. For cell type annotation with spatial transcriptomics, the raw count matrix was aligned to a constructed single cell reference including tumor cells, immune cells^83^ and stromal cells^84,85^ using the spacexr package (v2.2.1) with the Robust Cell Type Decomposition (RCTD) algorithm^86^. The RCTD doublet mode was used to assign a cell type to each bin 50. The RCTD full mode was applied to calculate the cell type fraction within each bin 50. For macrophage subtype annotation, the expression of macrophage cells were normalized using SCTransform (v2) to remove technical variability including sequencing depth. Macrophage subtypes were identified using the FindNeighbors function with the top 30 dimensions, followed by the FindClusters function with a resolution of 0.2. The top 100 feature genes for each macrophage subtype were listed in Supplementary Table 7. For spatial neighborhood analysis, each cell type neighbor ratio was qualified using the following definition:

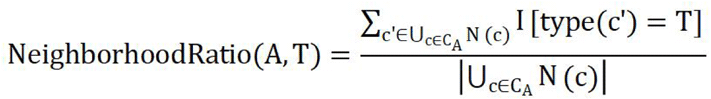

For cell type A, let *C_A_* = {c₁, c₂, …, cₙ } denote the complete set of all type A cells. For each cell c ∈ *C_A_*, its 10 cell neighborhood is defined as N(c) = {c’₁, c’₂, …, c’₁₀ }, where T represents a neighboring cell type of A. Cell neighbors identified using Spateo (v1.1.0)^87^ neighbors function. For spatial cell-cell communication analysis, CellChat (v2.1.2)^88^, with parameters tailored for spatial transcriptomics to restrict ligand–receptor interactions to short-range distances, was used to infer communications between tumor cell states and TME.

### Lineage tracing barcode analysis

For the analysis of target site editing efficiency of EP-eTRACER and NIH/3T3-eTRACER cells, we used the CRISPResso2 package (v2.2.13)^89^ to align reads to the reference sequence and identify mutation events. Reads with less than 70% alignment to the reference were excluded from qualitative analysis, and the edit event quantification window was defined as 30 bp flanking the classical Cas9 cleavage site. The edit ratio, edit size and edit diversity were quantified for each target gene. The edit ratio was calculated as the proportion of edited reads relative to total sequencing reads. Only insertion and deletion were included in the analysis of edit sizes. The correlation between the number of unique edits and the number of edited reads was calculated to evaluate edit diversity.

Lineage tracing barcodes, including static and evolving barcodes, derived from scRNA-seq, scMultiome-seq and Stereo-seq data were used for tumor clone identification and cell phylogeny reconstruction. To identify high quality lineage tracing barcodes, cell barcodes and UMI barcodes, the Cassiopeia^90^ pipeline (v2.0.0) was used for read quality control, error correction and noise removal. Single-cell lineage barcode sequencing reads from three stages, with a default minimum PHRED sequencing quality score of 10, were retained. Cell barcodes were filtered using platform-specific whitelist files, including 3M-february-2018.txt for scRNA-seq data and 737K-arc-v1.txt for scMultiome-seq, both available in the CellRanger software directory. For Stereo-seq data, reads were filtered based on spatial location barcodes. Only barcodes present in the corresponding barcodeToPos.h5 whitelist file were retained. Additionally, spatial location barcodes were error-corrected using ST_BarcodeMap (https://github.com/STOmics/ST_BarcodeMap) with the following parameters: barcodeStart = 36, barcodeLen = 25, umiStart = 76, umiLen = 10, and allowing up to 1 bp mismatch in the spatial barcodes. Reads with the same UMIs were collapsed into consensus sequences, allowing a maximum of 3 mismatches and 2 indels between the sequences of two aligned segments. The consensus sequence with the greatest number of UMIs was selected to represent the unique transcript of the lineage barcode. For the target site barcodes, sequences were aligned to the reference target site sequences using the Smith-Waterman local alignment algorithm, and edit types were identified within 30 bp on either side of the Cas9 cut site.

For clone detection of scRNA-seq data, the cells with the same unique static barcode combination are considered to belong to the same clone. To reduce noise from sequencing chimeras and cell-free transcripts in emulsion droplets, each barcode transcript must have a minimum of 10 sequencing reads, and each barcode must be supported by at least 2 distinct transcripts. Barcodes with a feature sequence and within the static barcode whitelist were retained. The cell-barcode UMI count matrix was generated and normalized, followed by clustering using the Cassiopeia assign lineage groups function to identify clones. A static barcode was considered in the clone set if it was present in more than 10% of cells within the clone. A cell was included in the clone if its barcode expression accounted for at least 25% of the total barcode expression within the clone set relative to other clones. Finally, 173 clones were identified ranging in size from > 4,500 (clone 1) to ∼30 cells (clone 173). 2,796 distinct clones were detected from 12,863 static barcodes at the BL, while 293 clones were from 1,350 static barcodes at the EE/LE. For clone detection of scMultiome data, 2,511 of the 2,568 cells had detectable static barcodes. By aligning these barcodes to scRNA-seq clone barcodes, clone IDs were assigned to the cells. For spatial clone detection, static barcodes were identified in a total of 26,603 bin 50s. To mitigate spatial sparsity, bin 50s with a diameter of 25 μm were used as a unit, as cells from the same clone are theoretically spatially proximate. Clone static barcodes identified from scRNA-seq data were used as references to assign clone identities to bin 50 static barcodes. If multiple clone identities were present within a bin 50, the identity supported by the greatest number of UMIs was assigned to that bin 50. Using this strategy, approximately 886 bin 50s at the BL, 1,3422 bin 50s at the EE, 7351 bin 50s containing clone barcodes were assigned to a clone.

For phylogenetic tree reconstruction, cell division history was built using CRISPR-Cas9 induced indels on 28 target sites of 14 genes (excluding *Acta1* due to its low expression in scRNA-seq data; see Figure S3C), with two alleles per gene theoretically. Due to cell-free transcripts in emulsion droplets, sequencing artifacts, and limited single-cell capture efficiency, certain target sites in individual cells inevitably exhibit noise edits or dropout events. To reduce the noise edits, reads (150 bp) with more than 0.1 (15 bp) mismatches and fewer than 0.7 (90 bp) matched were excluded, the edits with greater than 50bp were filtered out. Each target transcript must have a minimum of 3 sequencing reads. Based on clonal editing events distribution, clones with fewer than six editing events were excluded from downstream analysis. For Stereo-seq data, bin 50 was used as the unit for spatial clone identification. Target transcripts within each bin 50 that shared the same edit type were aggregated. Each editing event is required to be supported by at least 10 sequencing reads. Only spatial edits matching the single-cell clone edit set are retained. Phylogenetic trees are reconstructed using bin 50s that contain at least 5 edits. CRISPR-induced edits occur randomly; thus, within a single cell, the two alleles typically exhibit distinct edit patterns. Clonal cells derived from a common ancestor share identical allele-specific edit types, provided that the same ancestral allele was edited. This characteristic ensures that cells originating from the same clone display consistent allele-specific edit profiles. The target-site edit types were converted into a binary matrix where: Let C = {c₁, c₂, …, cₙ } be the set of n single cells belonging to a tumor clone, let M = {m₁, m₂, …, mₚ } be the set of distinct edits detected across the clone. The binary edit matrix was defined as D ∈ {0,1}^nxp^ where:

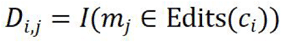

This results in a combination of allele-specific edits for each cell. Pairwise cell edit distances are computed using the Hamming distance:

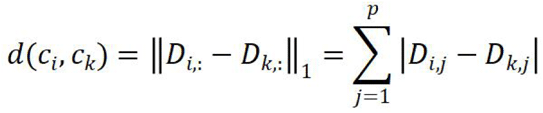

IQ-TREE2 (v2.2.2.7)^42^ was used to perform maximum-likelihood inference based on binary edit matrix. The polytomy parameter was used for cells without edit distance. The root of the tree was set as a synthetic barcode with no mutation (wild type). To evaluate the robustness of the phylogenetic trees, bootstrap method was used with 1,000 rounds of ultrafast bootstrap approximation and 1,000 rounds of SH-like approximate likelihood ratio test. The tree branching index is calculated as the number of internal branching nodes divided by (the number of terminal nodes−1). For phylogenetic tree visualization, ggtree (v3.10.1)^91^ was used with no branch lengths displayed.

For tumor cell plasticity quantification, the Fitch-Hartigan maximum-parsimony algorithm was used to infer the minimum number of times that tumor cell states had to have changed to give rise to the observed pattern. This function is implemented in the Cassiopeia package. The resulting score ranges between 0 and 1. A score of 0 indicates no changes to another state at all, while a score of 1 indicates that every cell has a different state from its sisters.

For cell-state transition probability inference, by coupling the cell phylogeny and their cell states, cell-state transition probabilities were quantified with PATH (v1.0)^43^. PATH adapts phylogenetic signal metric Moran’s *I*, which measures the degree to which related cells phenotypically resemble each other, to define a metric of the heritability versus plasticity of cellular phenotypes. The cell state transition dynamics can be linked to phylogenetic correlations by modeling state evolution as a Markov process along the phylogeny. While phylogenetic cell state pair frequencies provide empirical summaries of lineage-dependent dependencies, transition dynamics are formally inferred through a model defined on the tree, from which phylogenetic correlations are subsequently derived. Cell-state transitions were visualized using Dynamo (v1.4.0)^92^.

## Data and code availability

All processed data generated in this study have been deposited and are available at Zenodo (https://zenodo.org/records/19230230)^93^. Raw data are publicly available from the National Genomics Data Center under the accession numbers PRJCA040871 (https://ngdc.cncb.ac.cn/gsa/s/fX633AQn). All computer code used in this study is available from the GitHub repositories (https://github.com/liangzhenhou/eTRACER).

## Acknowledgements

The authors thank the members of Ji and Hu lab for constructive discussions. This work was supported by the National Key Research and Development Program of China (2022YFA1103900 and 2020YFA0803300 to H.J.), the National Natural Science Foundation of China (32525022 and 82241236 to Z.H., 82341002, 32293192 and 82030083 to H.J., 82372763 to X.W., 82273400 to Y.J.), the Innovative research team of high-level local universities in Shanghai (SSMU-ZLCX20180500 to H.J.).

## Author contributions

H.J., Z.H., X.W. and J.Y. conceived and designed the study. J.Y., X.W. and N.Z. performed molecular, cell culture and mouse experiments. L.H., Z.L., D.X. K.W., L.T.O.L. and X.H. analyzed the sequencing data. Y.B. participated in the creation of the schematic illustration. Y.F. offered the technical support for FACS. Y.X.C., R.W., Y.J., Y.C., X.C., J.X.H. and X.T. participated in molecular experiments. H.J., Z.H., J.Y., X.W., and L.H. wrote the manuscript. H.J. and Z.H. supervised the project.

---

**Figure S1.**
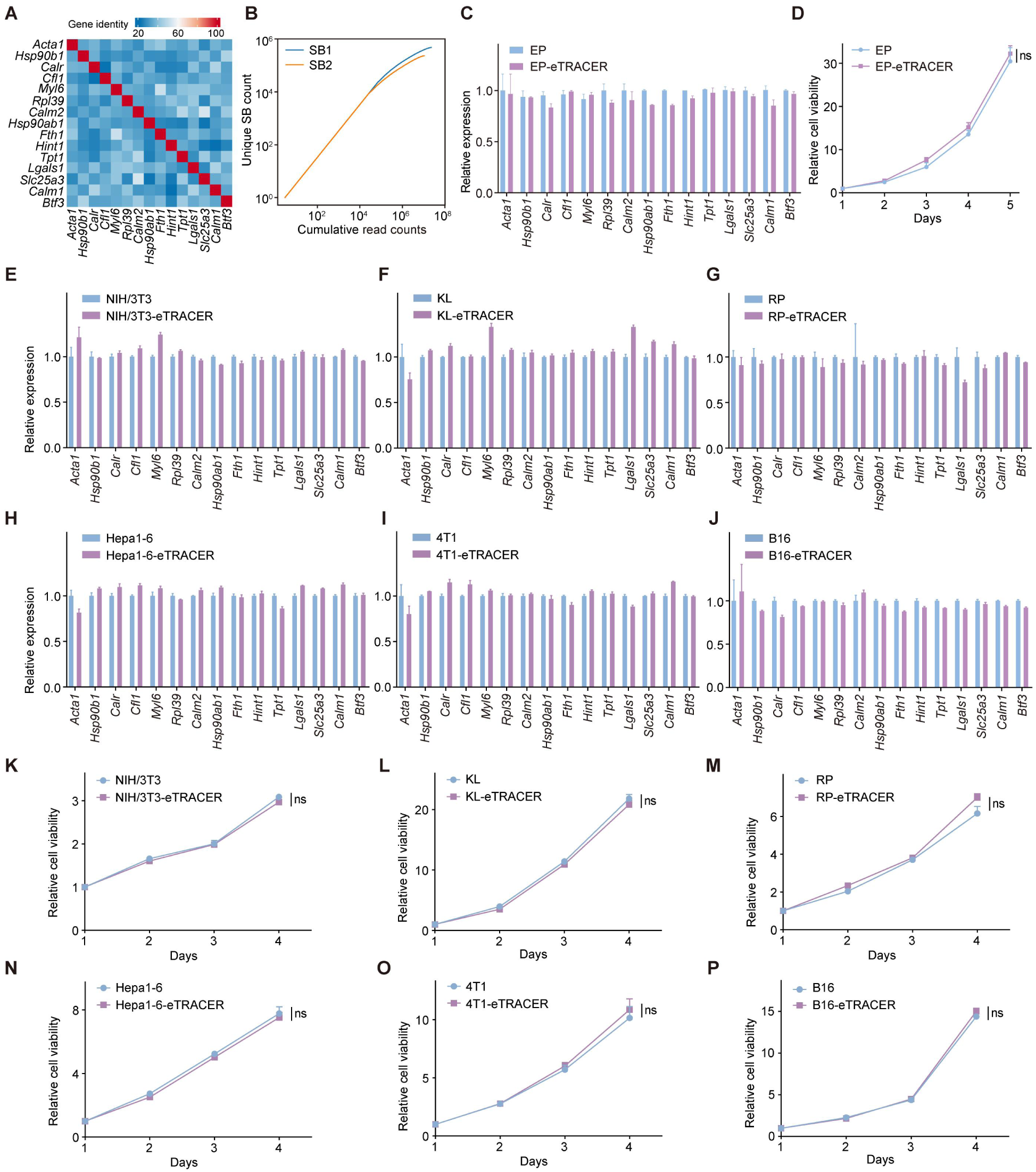
*In vitro* validation of eTRACER. (A) DNA sequence similarity among the target sites in the 3′ UTRs of 15 endogenous genes was analyzed using multiple sequence alignment. (B) The relationship between the number of unique static barcodes (SB) and cumulative sequencing reads. SB1 and SB2 contain 474,796 and 232,094 unique static barcodes, respectively. (C) Expression of individual endogenous genes in EP (*EGFR^L858R^;Trp53^-/-^*) and EP-eTRACER cells in the presence of Dox (0.5 µg/mL) for 2 weeks assessed by qRT-PCR. Mean±s.e.m. (D) Viability of EP and EP-eTRACER cells cultured with Dox for 2 weeks. Two-sided Student’s t-test, mean±s.e.m. (E-P) The expression of 15 endogenous genes as well as cell viability were examined in mouse parental cells and eTRACER cells in the presence of Dox (0.5 µg/mL) for 2 weeks. NIH/3T3 (E,K), *Kras^G12D^;Lkb1^-/-^* (KL, lung adenocarcinoma) (F,L), *Rb1^-/-^;Trp53^-/-^* (RP, small cell lung cancer) (G,M), Hepa1-6 (hepatocellular carcinoma) (H,N), 4T1 (breast cancer) (I,O) and B16 (melanoma) (J,P). Two-sided Student’s t-test, \**P* < 0.05; \*\**P* < 0.01; \*\*\**P* < 0.001; \*\*\*\**P* < 0.0001; ns, not significant, *P* ≥ 0.05. Mean±s.e.m.

**Figure S2.**
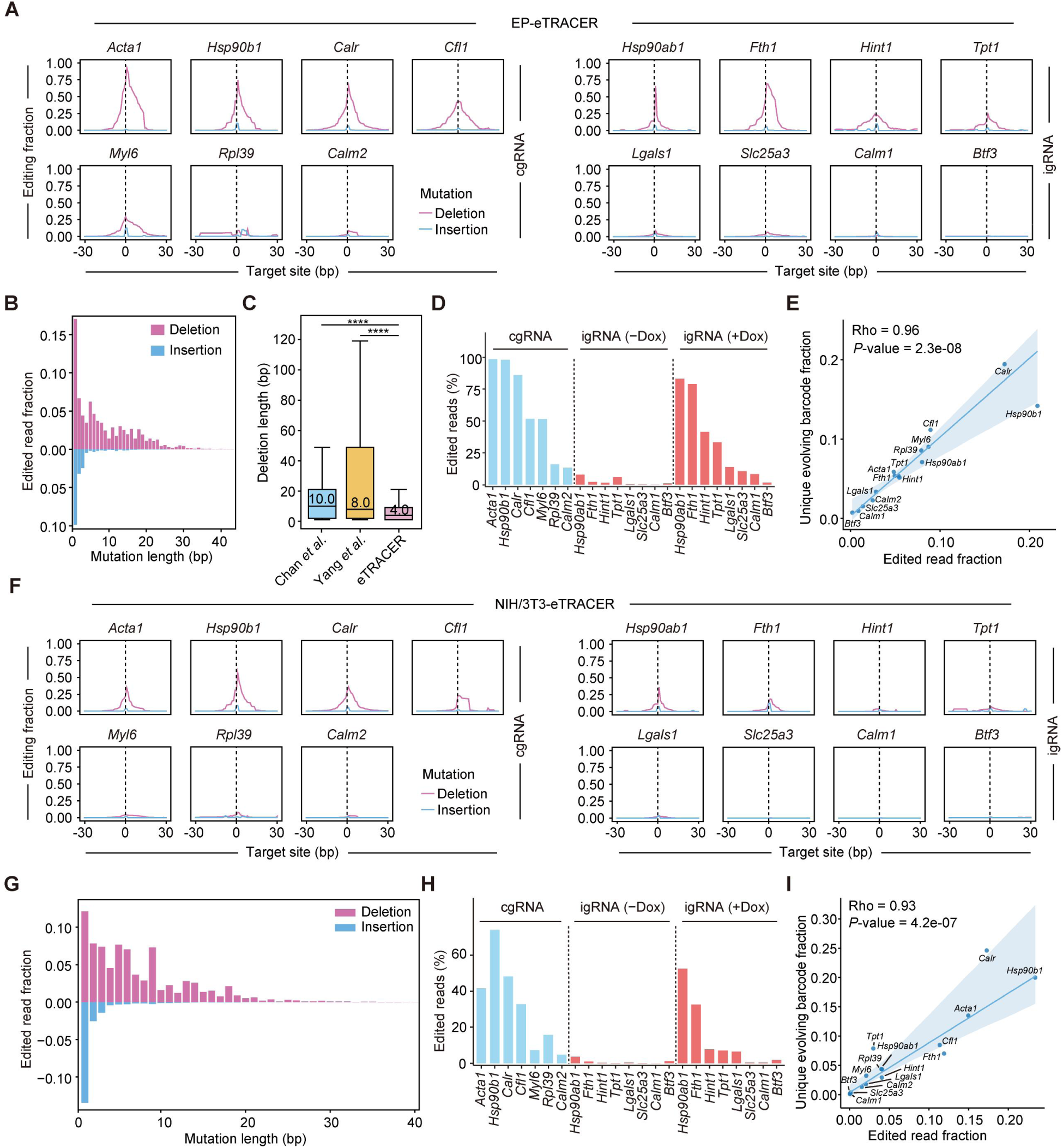
*In vitro* target editing efficiency of eTRACER. (A) The CRISPR editing profiles of endogenous target sites in EP-eTRACER cells. Dashed line denoted the Cas9 cleavage site. (B) Distribution of insertion/deletion (indel) lengths across all endogenous sites in EP-eTRACER cells. (C) Comparison of deletion lengths induced by eTRACER and CRISPR-Cas9 tracers in previous studies (Chan *et al.*^23^; Yang *et al.*^21^). The values shown indicate the median deletion length (in bp). Wilcoxon test, \**P* < 0.05; \*\**P* < 0.01; \*\*\**P* < 0.001; \*\*\*\**P* < 0.0001; ns, not significant, *P* ≥ 0.05. (D) Editing efficiencies of individual endogenous sites in EP-eTRACER cells following 2-week culture with or without Dox. (E) Correlation between unique evolving barcodes and edited reads on individual endogenous sites in EP-eTRACER cells following 2-week culture with Dox. Spearman’s rho and *P-*value are shown. (F) The CRISPR editing profiles of endogenous target sites in NIH/3T3-eTRACER cells. Dashed line denoted the Cas9 cleavage site. (G) Distribution of indel lengths across all endogenous target sites in NIH/3T3-eTRACER cells. (H) Editing efficiencies of endogenous target sites in NIH/3T3-eTRACER cells following 2-week culture with or without Dox. (I) Correlation between the unique evolving barcodes and edited reads from individual target sites in NIH/3T3-eTRACER cells. Spearman’s rho and *P-*value are shown.

**Figure S3.**
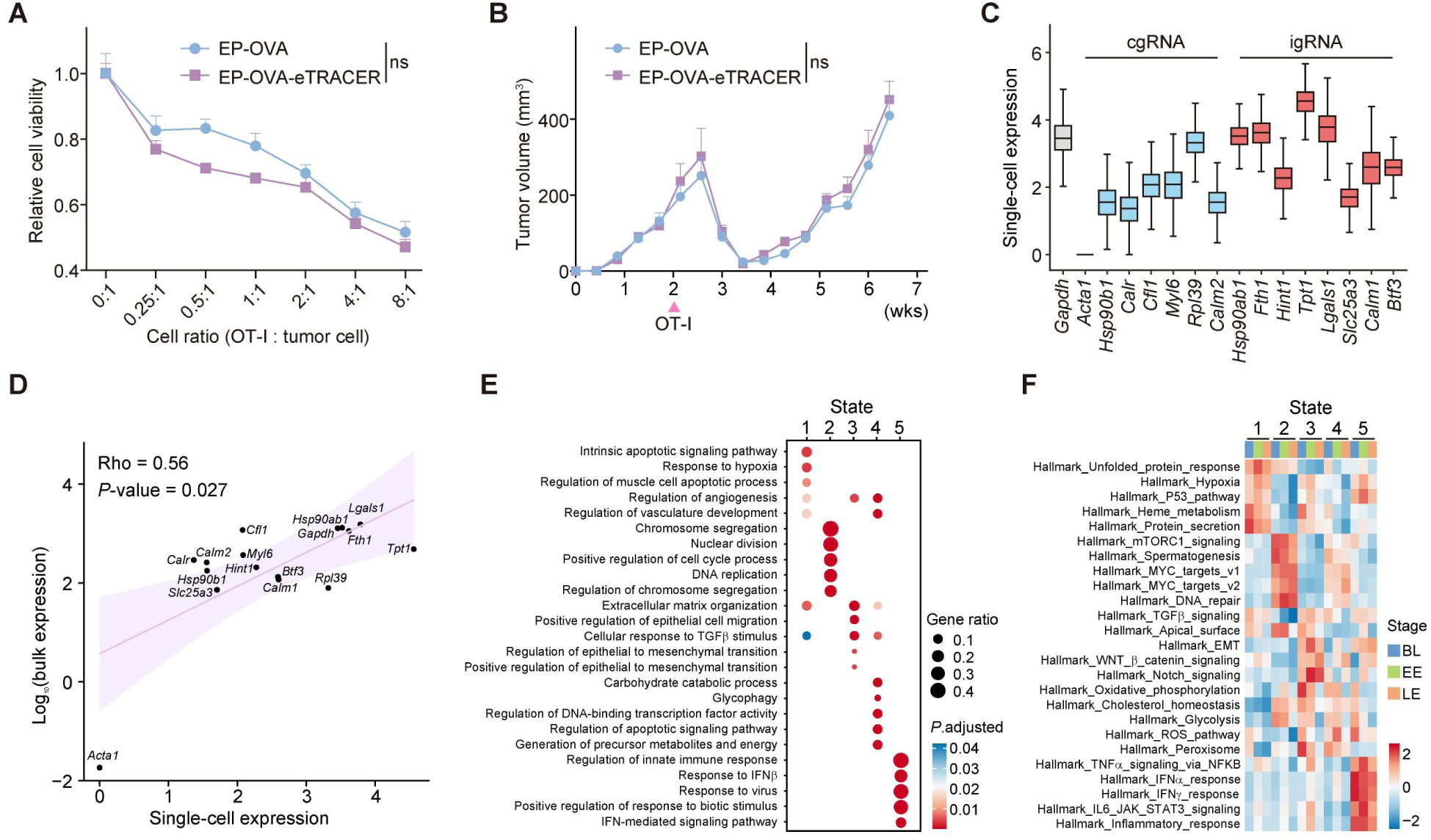
eTRACER validation and pathway enrichment analysis across cell states. (A) Co-culture cytotoxicity assay evaluating the response of EP-OVA and EP-OVA-eTRACER to OT-I cell-mediated cytotoxicity. Two-sided Student’s t-test. Mean±s.e.m. (B) Tumor growth of EP-OVA and EP-OVA-eTRACER in *Rag2^-/-^* mice following OT-I cell treatment. At 1 week post-engraftment of 50,000 tumor cells, Dox was administered to mice in drinking water (1 mg/mL). Two-sided Student’s t-test. Mean±s.e.m. (C) Expression of 15 endogenous genes based on scRNA-seq data analyses. (D) Correlation of individual endogenous gene expression between scRNA-seq and bulk RNA-seq data. Spearman’s rho and *P-*value are shown. (E-F) Gene Ontology (GO) (E) and hallmark analyses (F) of signaling pathways enriched in individual tumor cell states.

**Figure S4.**
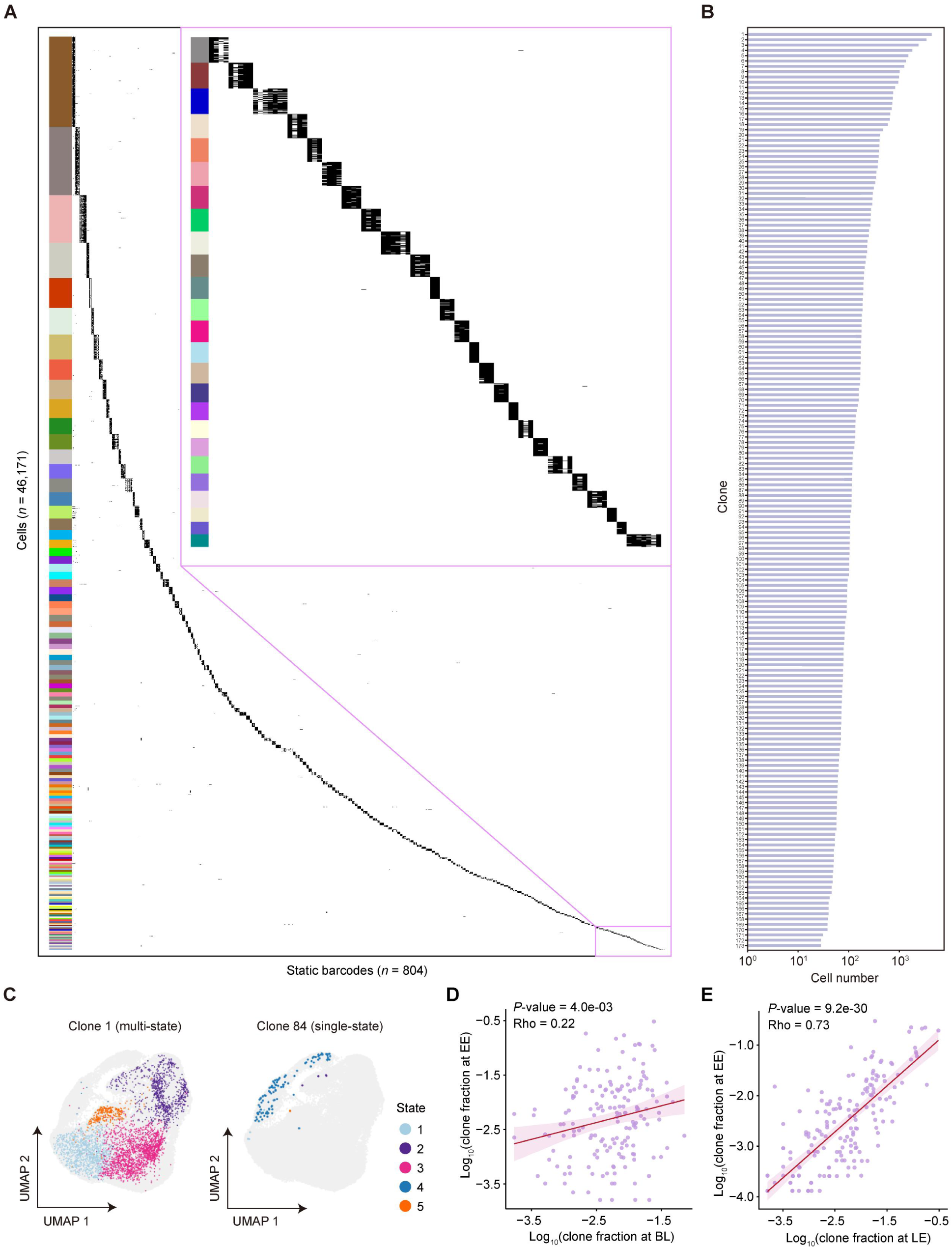
Clone identification and clonal dynamics analysis. (A) The unique static barcodes (rows, n = 804) among the 46,171 cells (columns) in top 173 clones with > 15 cells. Each color represents a distinct clone, with clones 1 to 173 arranged sequentially from top to bottom. (B) The individual cell numbers of top 173 clones. (C) UMAP visualization of a representative multi-state clone (clone 1) and a single-state clone (clone 84). (D-E) Comparative analyses of clone fractions between BL and EE stages (D) and between EE and LE stages (E). Spearman’s rho and *P-*value are shown.

**Figure S5.**
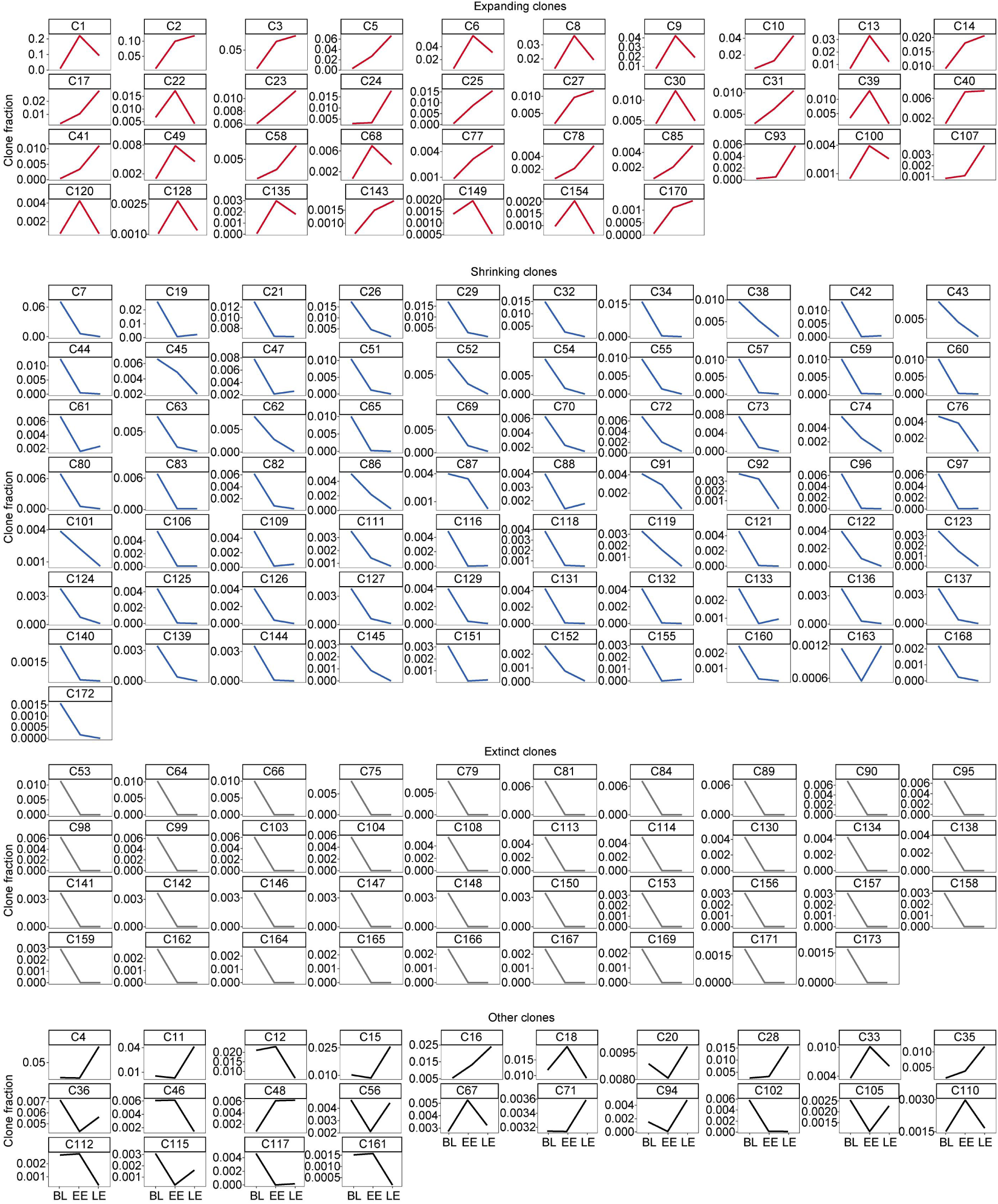
Clonal growth dynamics across the three stages, BL, EE and LE. Line plots showing the proportion changes of individual clones across the three stages, grouped by clone fates. The top 173 clones (each > 15 cells) are shown.

**Figure S6.**
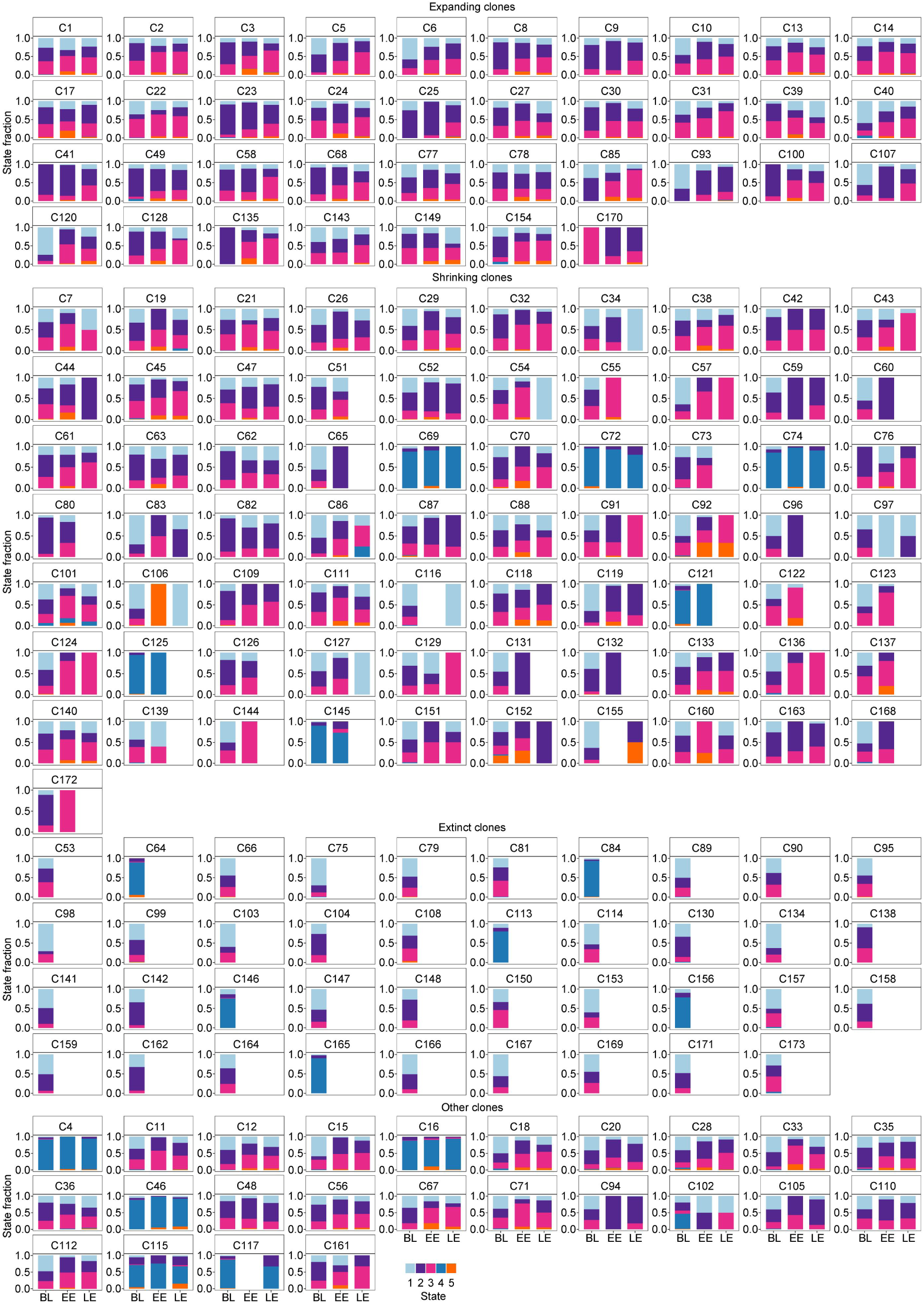
Cell state compositions within clones across three fate groups. The top 173 clones (each > 15 cells) are shown. The number above the bar chart indicates the clone ID.

**Figure S7.**
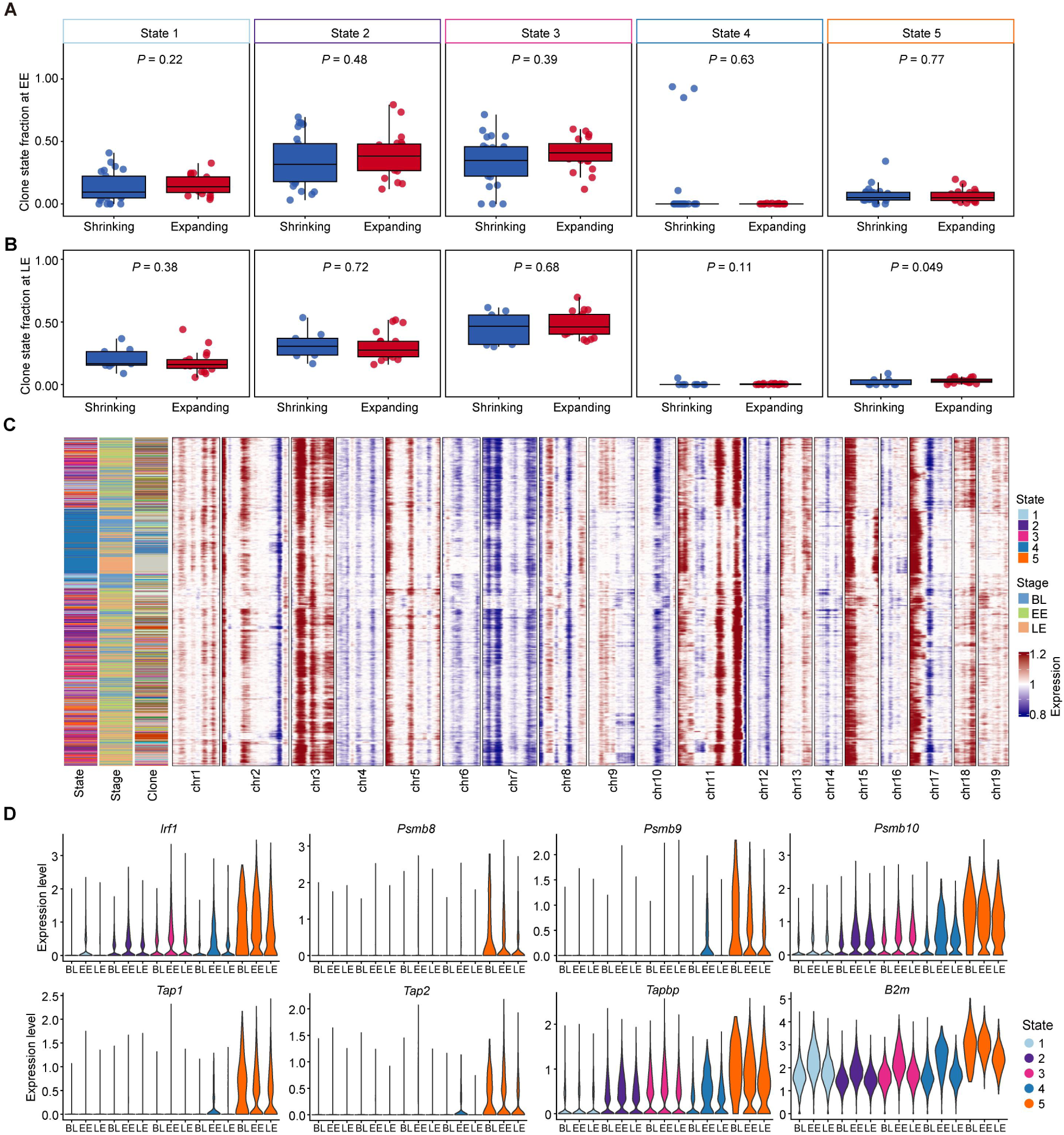
State compositions, CNV profiles, and antigen presentation gene expression. (A-B) Association between the proportion of each cell state in the EE (A) and LE (B) clones and clonal fate groups, assessed by the Kruskal-Wallis test. (C) CNV inference was performed using scRNA-seq data. The cells (rows) were ordered according to hierarchical clustering. For each state, 1,500 cells were randomly chosen for downsampling. (D) Expression profiles of representative antigen presentation genes across various stages and states.

**Figure S8.**
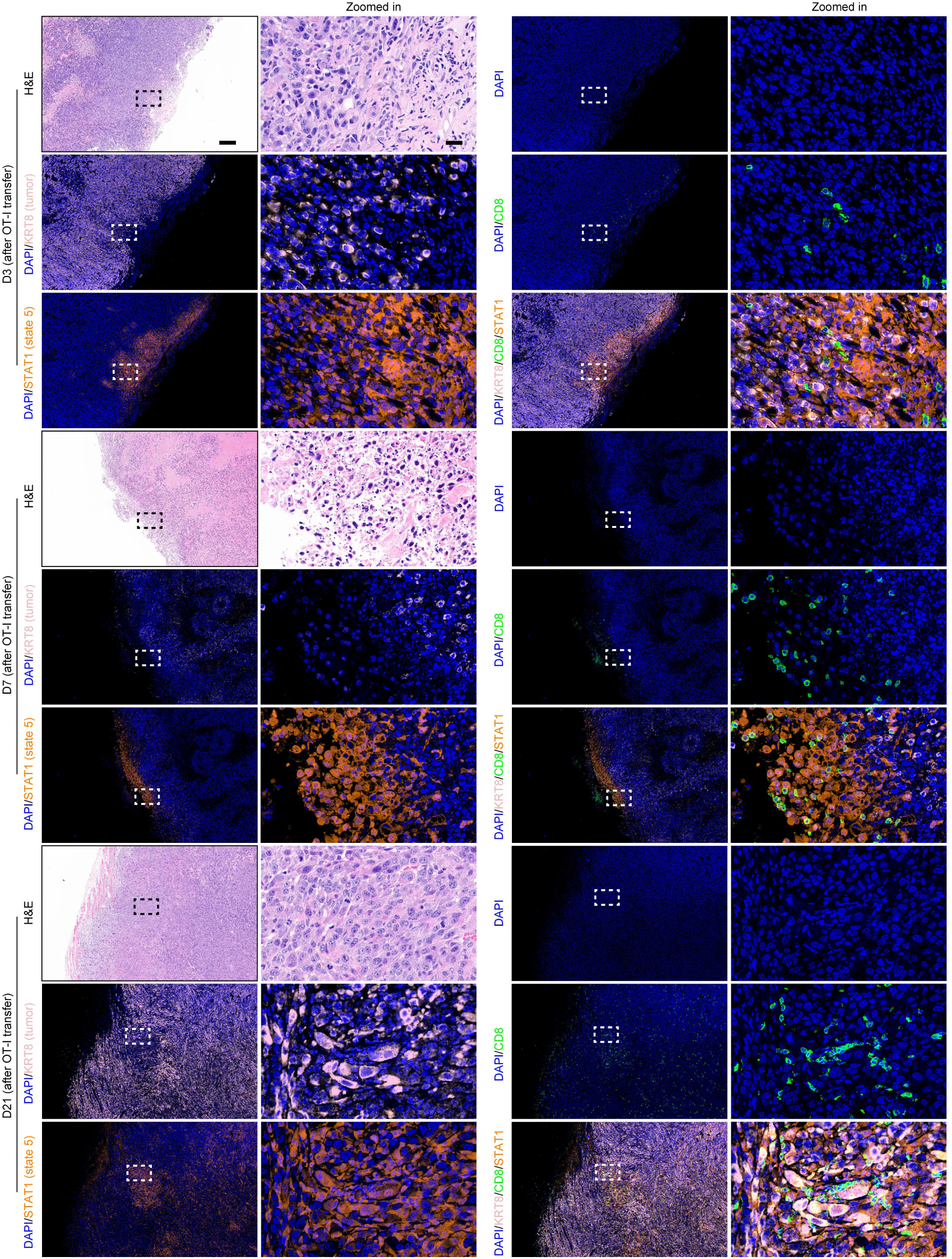
Spatial co-localization analysis of state 5 cells and OT-I cells. Representative H&E and immunofluorescence images showing KRT8, STAT1 and CD8 expression in EP-OVA-shCtrl tumor samples at day 3, 7 and 21 post-OT-I injection (see Figure S19A). Scale bars represent 200 µm and 20 µm (zoomed-in view), respectively.

**Figure S9.**
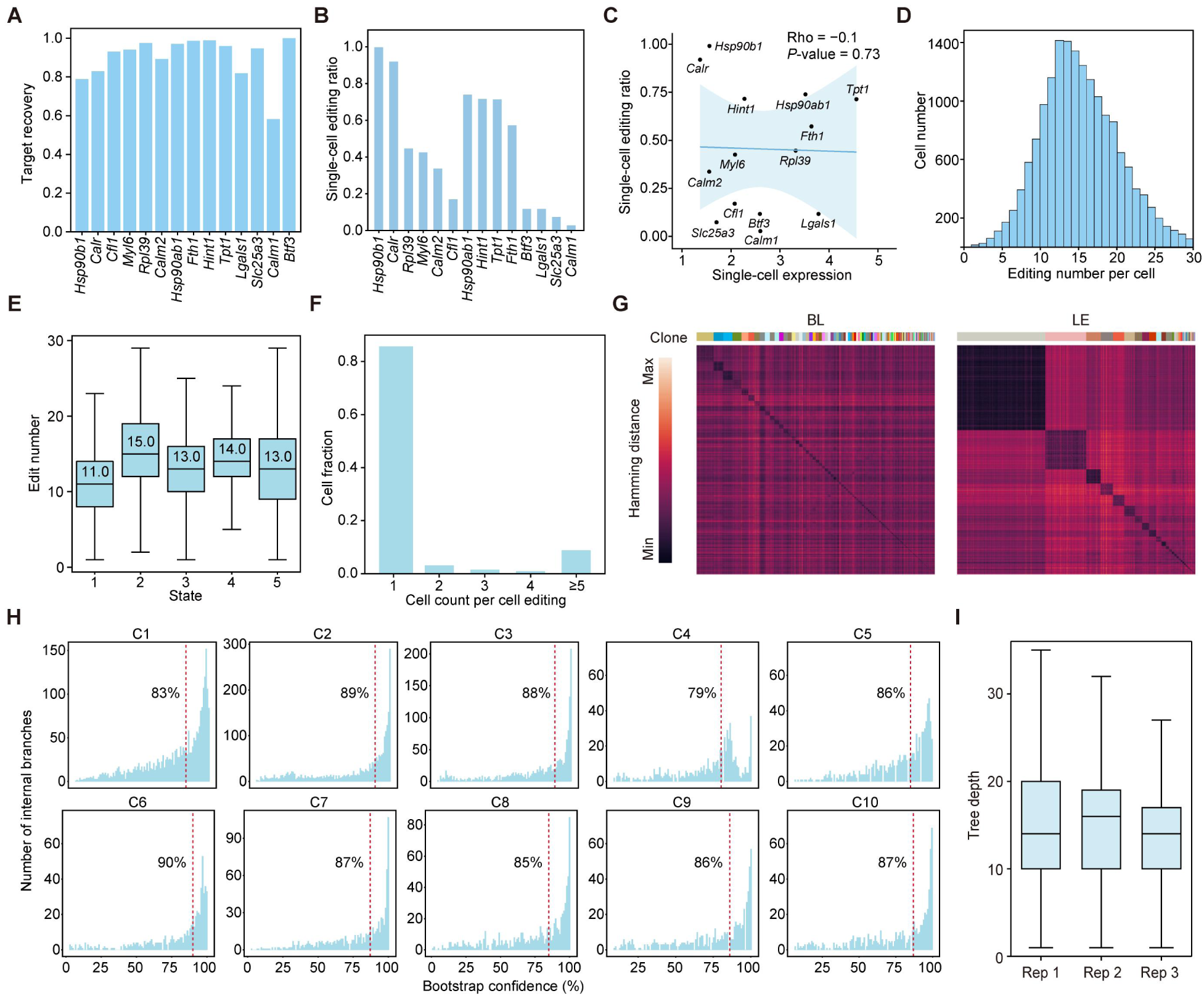
eTRACER enables the reconstruction of high-fidelity single-cell phylogenies. (A) Recovery rates of endogenous target sites across 91,411 single cells. *Acta1* in EP cells was excluded due to low expression. (B) Editing efficiencies of target sites in 91,411 single cells. (C) Correlation between gene expression and editing efficiencies of target sites at single-cell level. Spearman’s rho and *P-*value are shown. (D) Distribution of cell number by the editing number per cell. The mean editing number per cell is 15. (E) Edit number across cell states. The median editing number per state is shown. (F) The fraction of uniquely barcoded cells (cell count = 1) and non-uniquely barcoded cells (cell count > 1) among 46,171 cells from the top 173 clones. (G) Heatmap showing edit distance of evolving barcodes within clones at the BL and LE stages. Representative data from replicate 2. (H) Bootstrapping values of the trees for the top 10 clones. Red dotted line indicated the median. (I) Statistical analyses of the phylogenetic tree depths in individual replicates.

**Figure S10.**
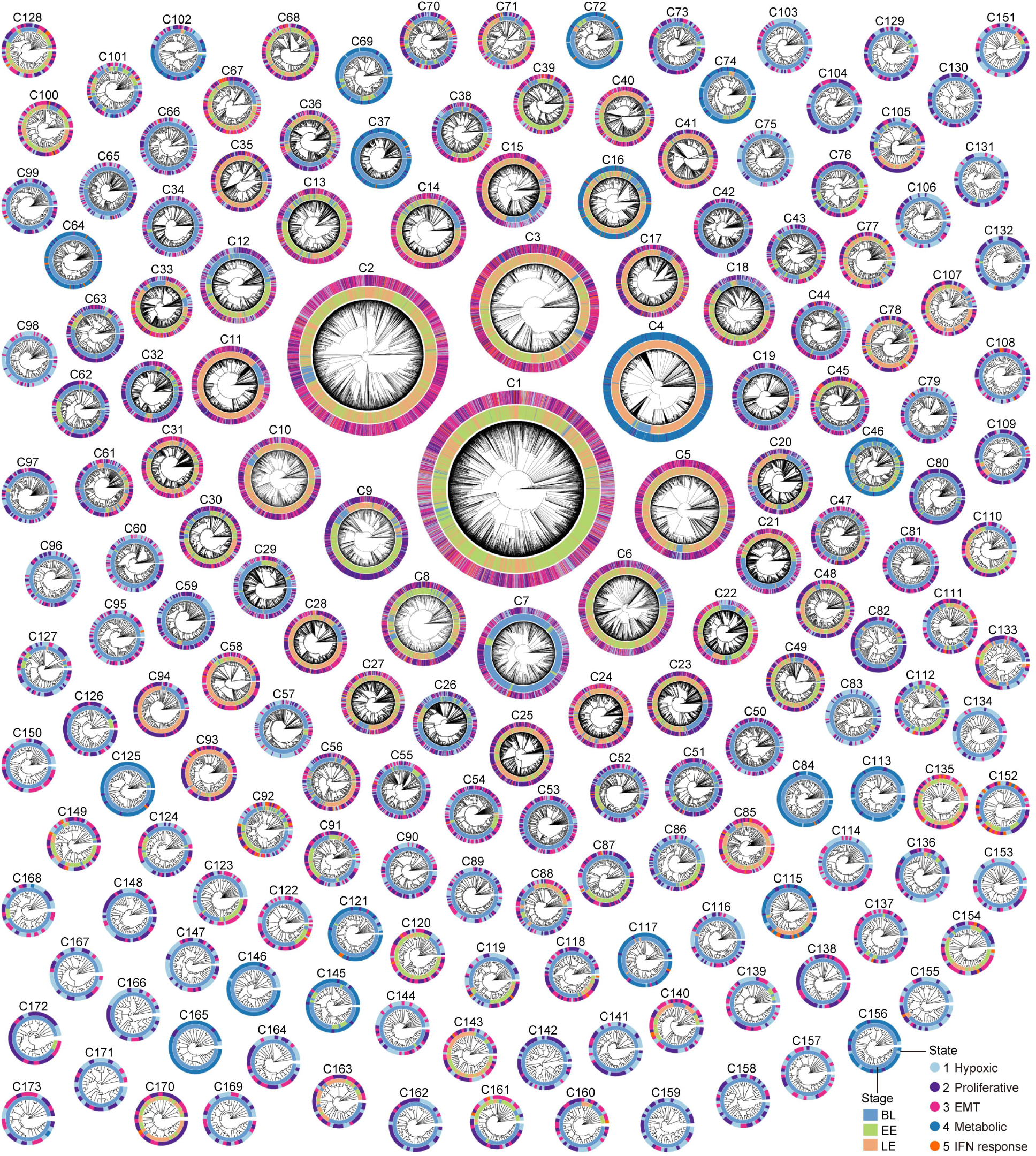
Multimodal lineage trees reconstructed by eTRACER. Single-cell phylogenetic trees for top 173 clones (each > 15 cells) are visualized as radial phylograms. Each cell is positioned along the circumference, colored according to its stage (BL, EE, LE) and transcriptomic state.

**Figure S11.**
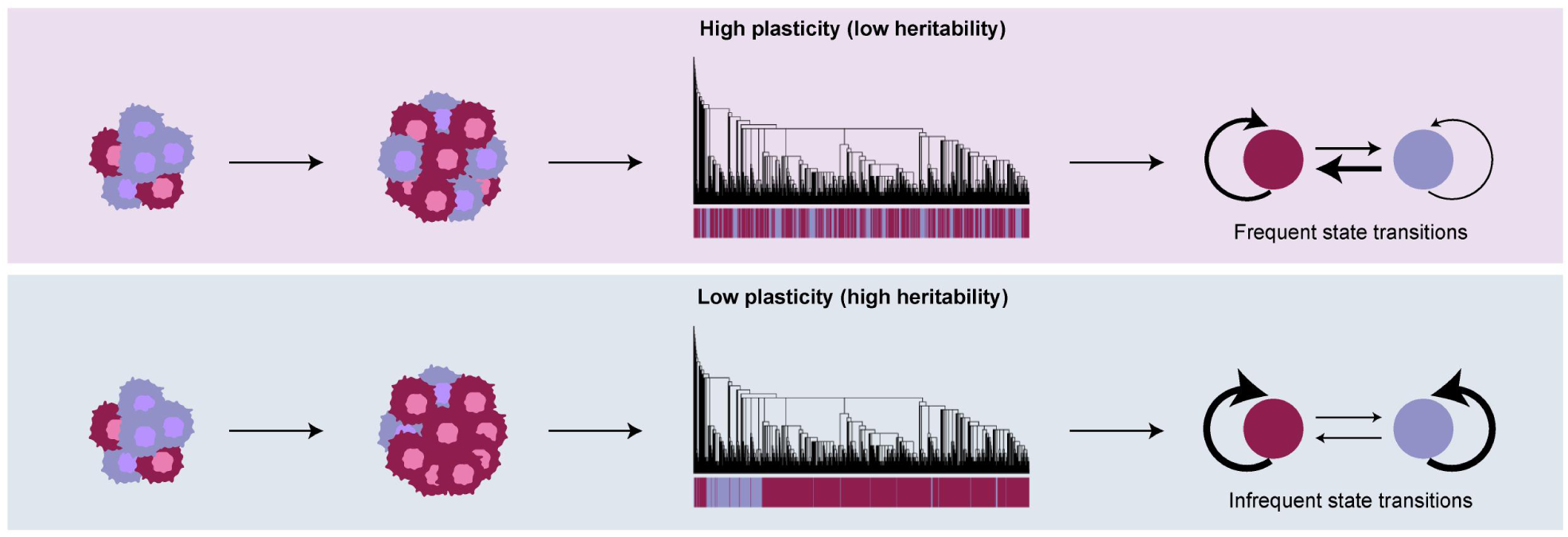
Clonal state plasticity model. High cell-state plasticity vs. low cell-state plasticity model.

**Figure S12.**
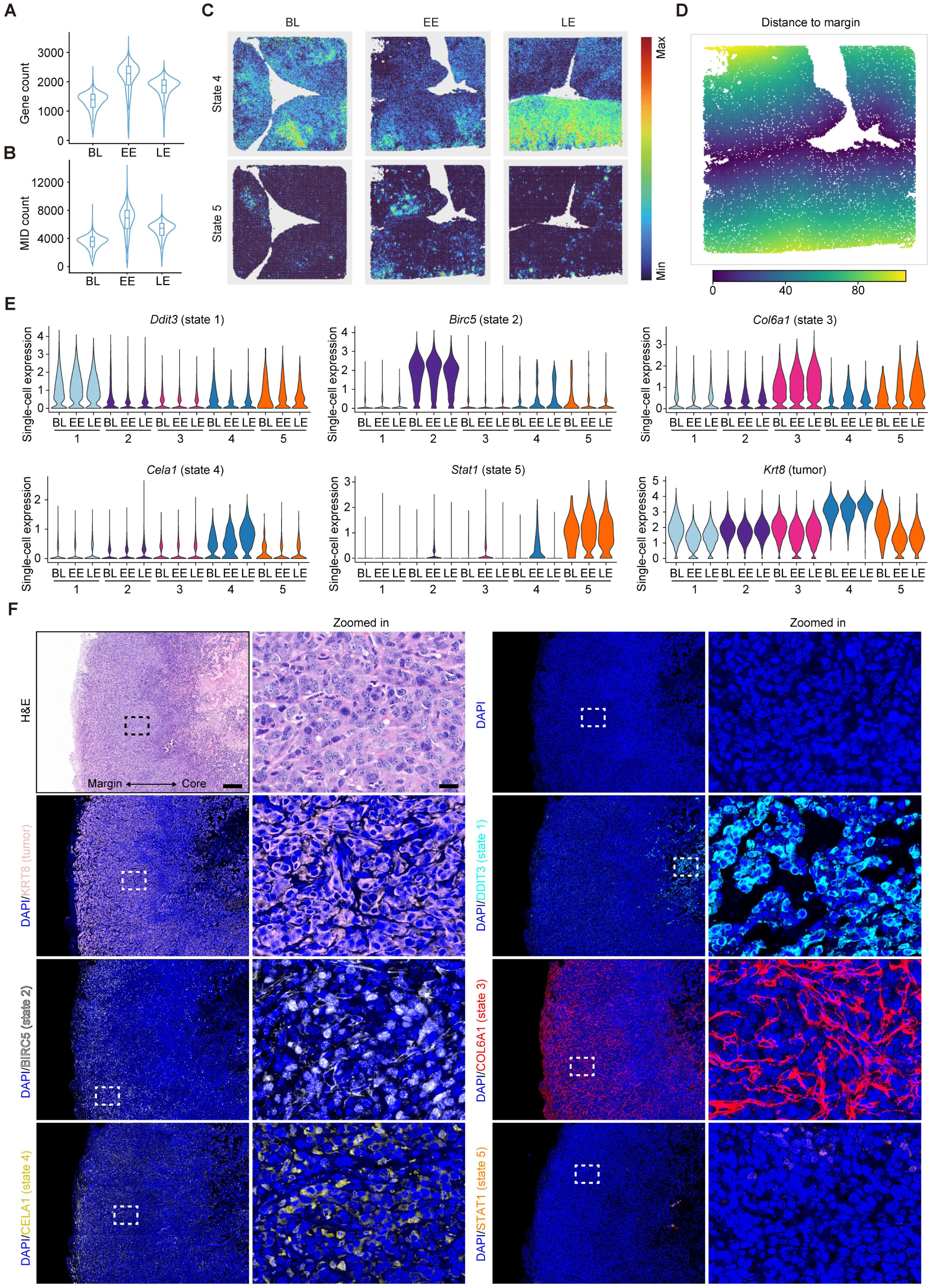
Spatial mapping of tumor cell states. (A-B) Gene counts (A) and Molecular IDentifier (MID) counts (B) at the BL, EE and LE stages based on spatial transcriptomic data analyses. (C) Spatial mapping of tumor cell state 4 and state 5 at the BL, EE and LE stages. (D) Visualization of cell distances to the tumor margin. (E) Expression of state-specific marker genes and the tumor marker *Krt8*. Numbers indicate cell states. (F) Representative H&E and immunofluorescence images showing KRT8, DDIT3, BIRC5, COL6A1, CELA1 and STAT1 expression in EP-OVA-shCtrl tumor samples at day 3 post-OT-I injection (see Figure S19A). Scale bars represent 200 µm and 20 µm (zoomed-in view), respectively.

**Figure S13.**
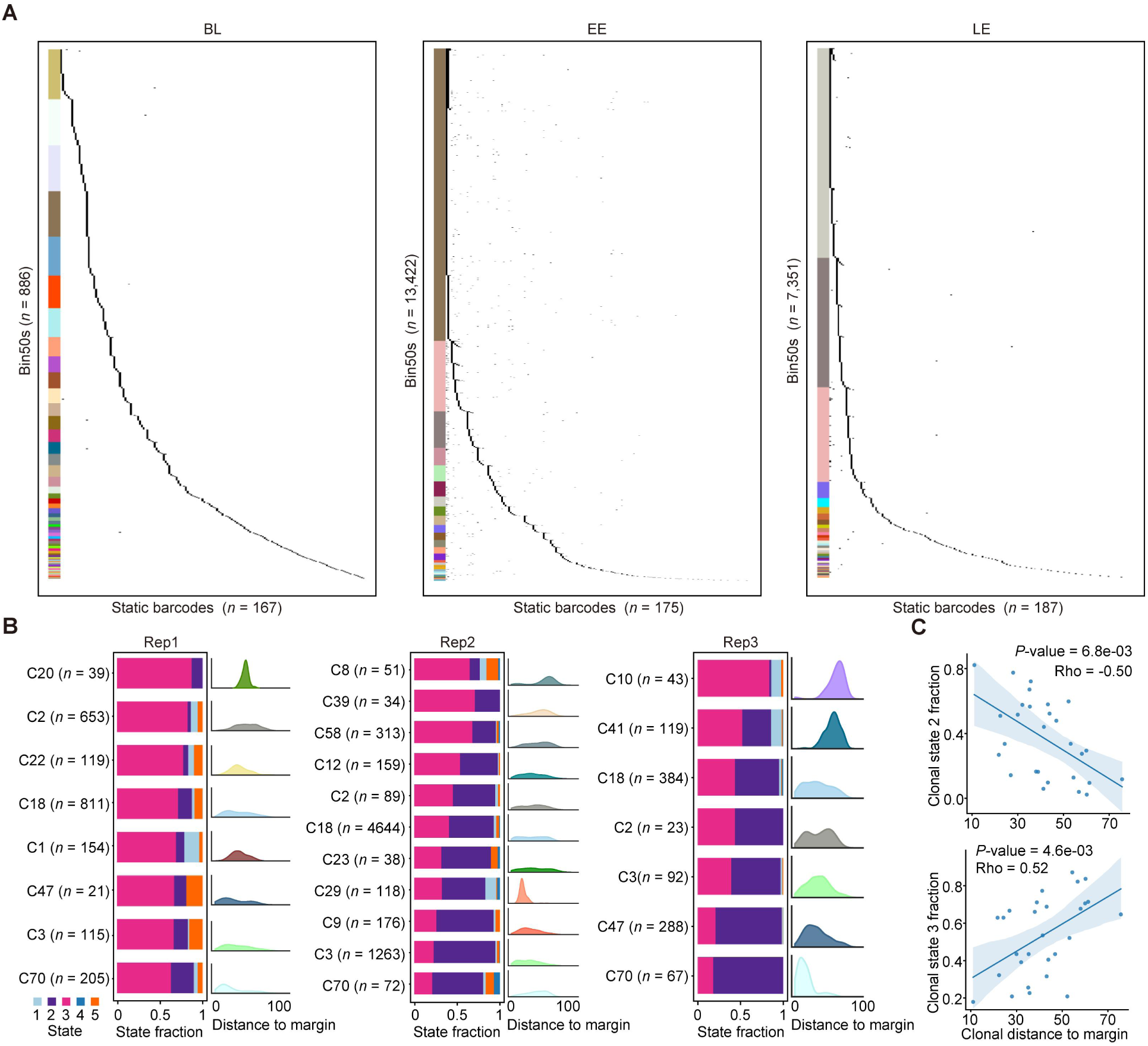
Spatial mapping of tumor clones. (A) The unique static barcodes (rows) among the bin 50s (columns) at the BL, EE, and LE stages. The unique color of each clone is consistent with the scRNA-seq data. (B) The state proportions of clones, along with density plots depicting their spatial distribution relative to the tumor margin at the EE stage. The unit of X-axis is bin 50 (25 μm). (C) The correlation between the proportion of state 2 or state 3 within tumor cell clones and their distance to tumor margin, respectively. Spearman’s rho and *P-*value are shown.

**Figure S14.**
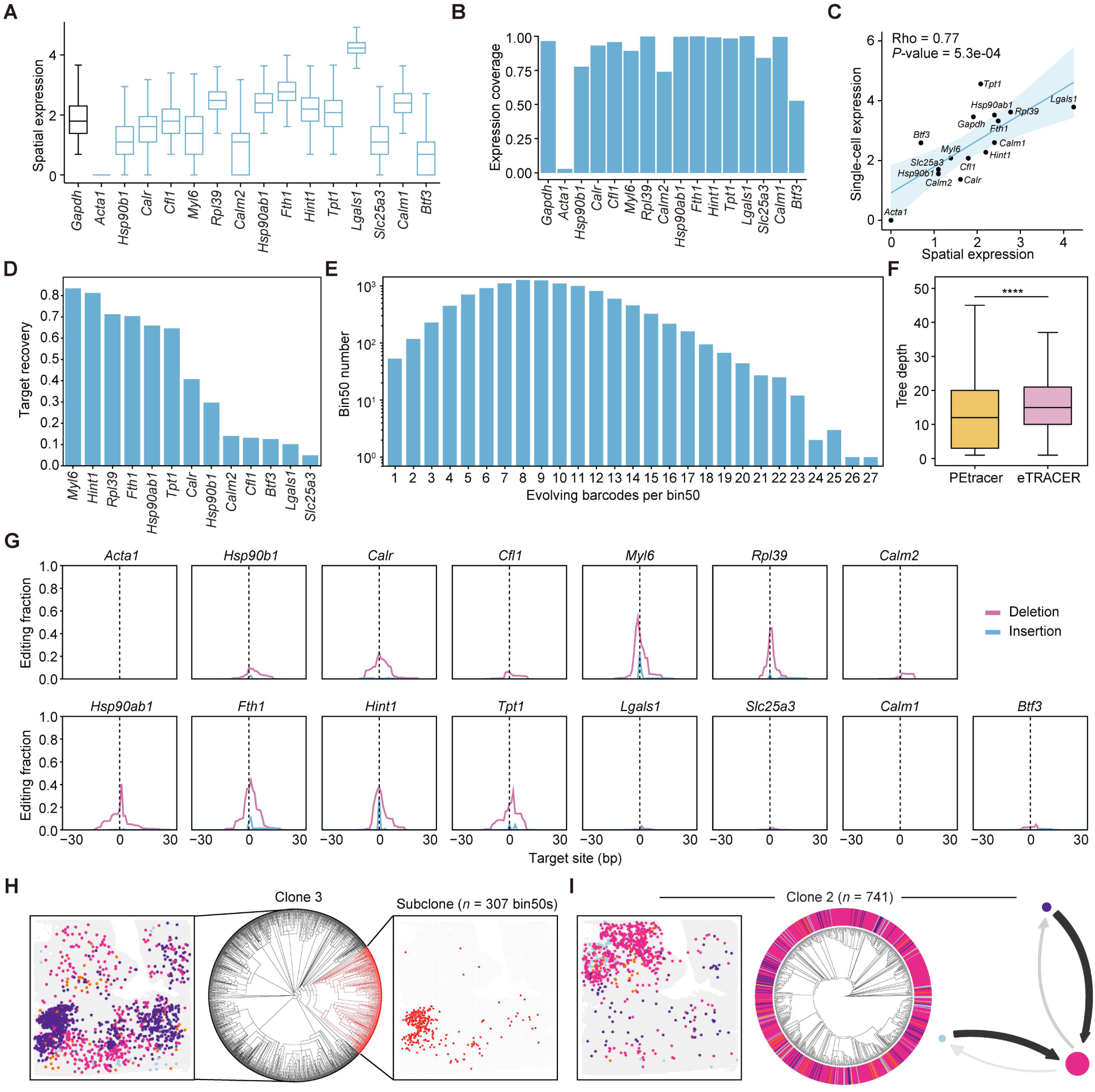
Robust spatial phylogenetic reconstruction with eTRACER. (A-B) Spatial transcriptomic expression levels (A) and capture efficiencies (B) of 15 endogenous genes. (C) Correlation of individual endogenous gene expressions between scRNA-seq and Stereo-seq data. Spearman’s rho and *P-*value are shown. (D) The recovery rate of bin 50s with specific edited target sites, excluding *Acta1* (undetectable expression) and *Calm1* (insufficient detection), at the EE stage based on Stereo-seq data. (E) The distribution of bin 50 counts with respect to the number of evolving barcodes per bin 50 at the EE stage. (F) The tree depth of eTRACER spatial lineage trees (n = 21 clones) is compared with that of PEtracer spatial lineage trees (n = 12 clones) from a previous study^31^. In both datasets, phylogenetic trees were constructed using the maximum likelihood method. Clones containing more than 2,000 cells were downsampled to 2,000 cells. Wilcoxon test. (G) The CRISPR editing profiles of individual target sites in tumor bin 50s at the EE stage. The dashed lines denote the Cas9 cleavage sites. (H) Spatial mapping and the phylogenetic tree of representative clone 3 at the EE stage. The subclone is highlighted in red on the phylogenetic tree, with its corresponding spatial distribution mapped. (I) Spatial locations, phylogenetic trees and state-transition networks of clone 2.

**Figure S15.**
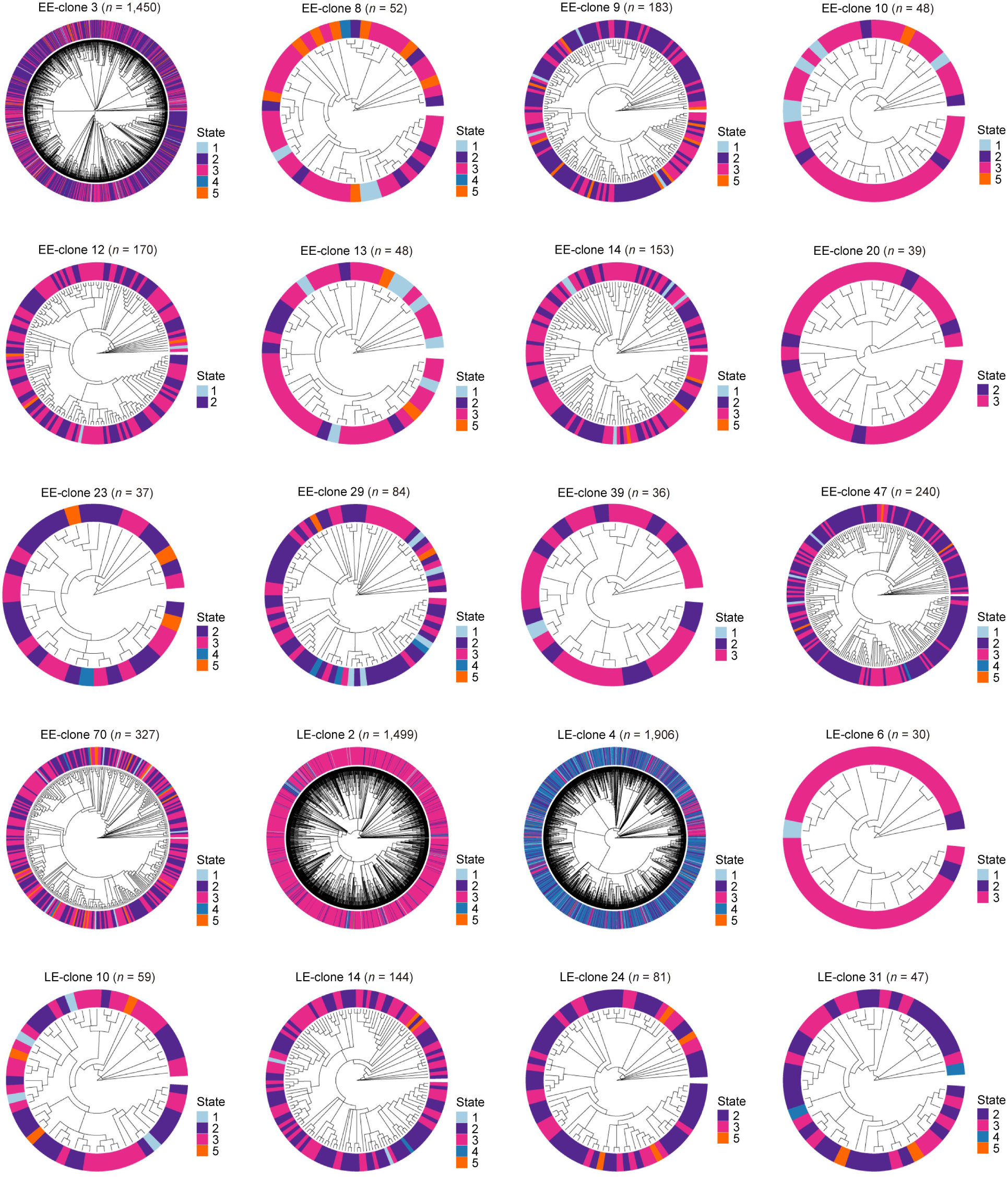
eTRACER enables high-resolution spatial phylogenetic reconstruction. Representative phylogeny reconstruction based on Stereo-seq data analyses.

**Figure S16.**
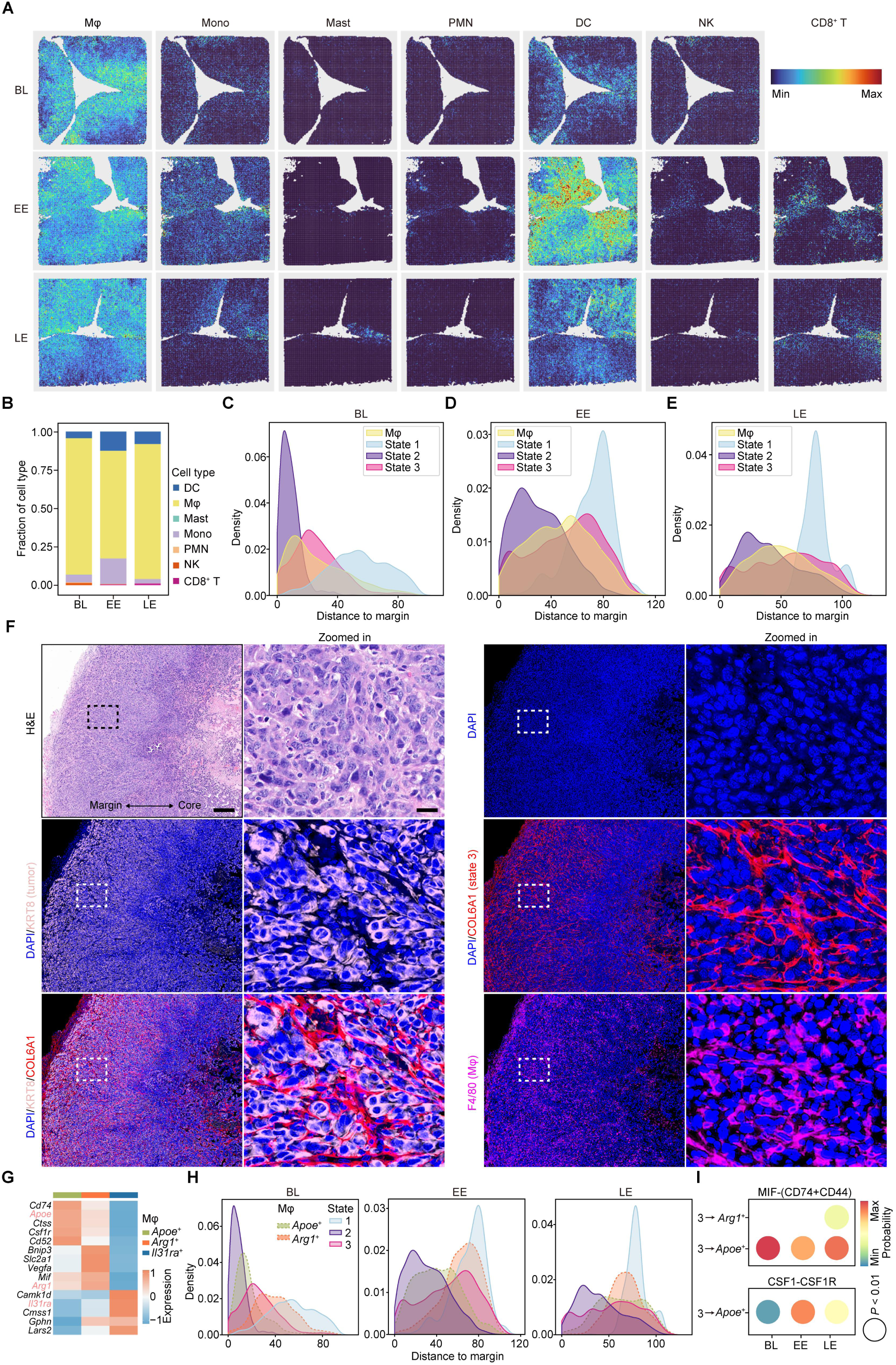
Spatial crosstalk of State 3 cells and macrophages underlying tumor adaptation. (A) Spatial mapping of immune cells at the BL, EE and LE stages. (B) Fractions of immune cells across BL, EE, and LE stages derived from spatial transcriptomic data. (C-E) Quantitative analyses of spatial distances to tumor margin for cell states and macrophages across BL (C), EE (D) and LE (E) stages. The area under the curve for each state summed up to 1. The unit of X-axis is bin 50 (25 μm). (F) Representative H&E and immunofluorescence images showing KRT8, COL6A1 and F4/80 expression in EP-OVA-shCtrl tumor samples at day 3 post-OT-I injection (see Figure S19A). Scale bars represent 200 µm and 20 µm (zoomed-in view), respectively. (G) Heatmap illustrating the marker gene expression in individual macrophage subtypes. (H) Quantitative analyses of spatial distances to tumor margin for tumor cell states and macrophage subtypes at the BL, EE and LE stages. The area under the curve for each state summed up to 1. The unit of X-axis is bin 50 (25 μm). (I) The quantitative probabilities of MIF-(CD74+CD44) and CSF1-CSF1R-mediated state 3 and macrophage cell interactions by cell-cell communication analyses. *P*-value < 0.01.

**Figure S17.**
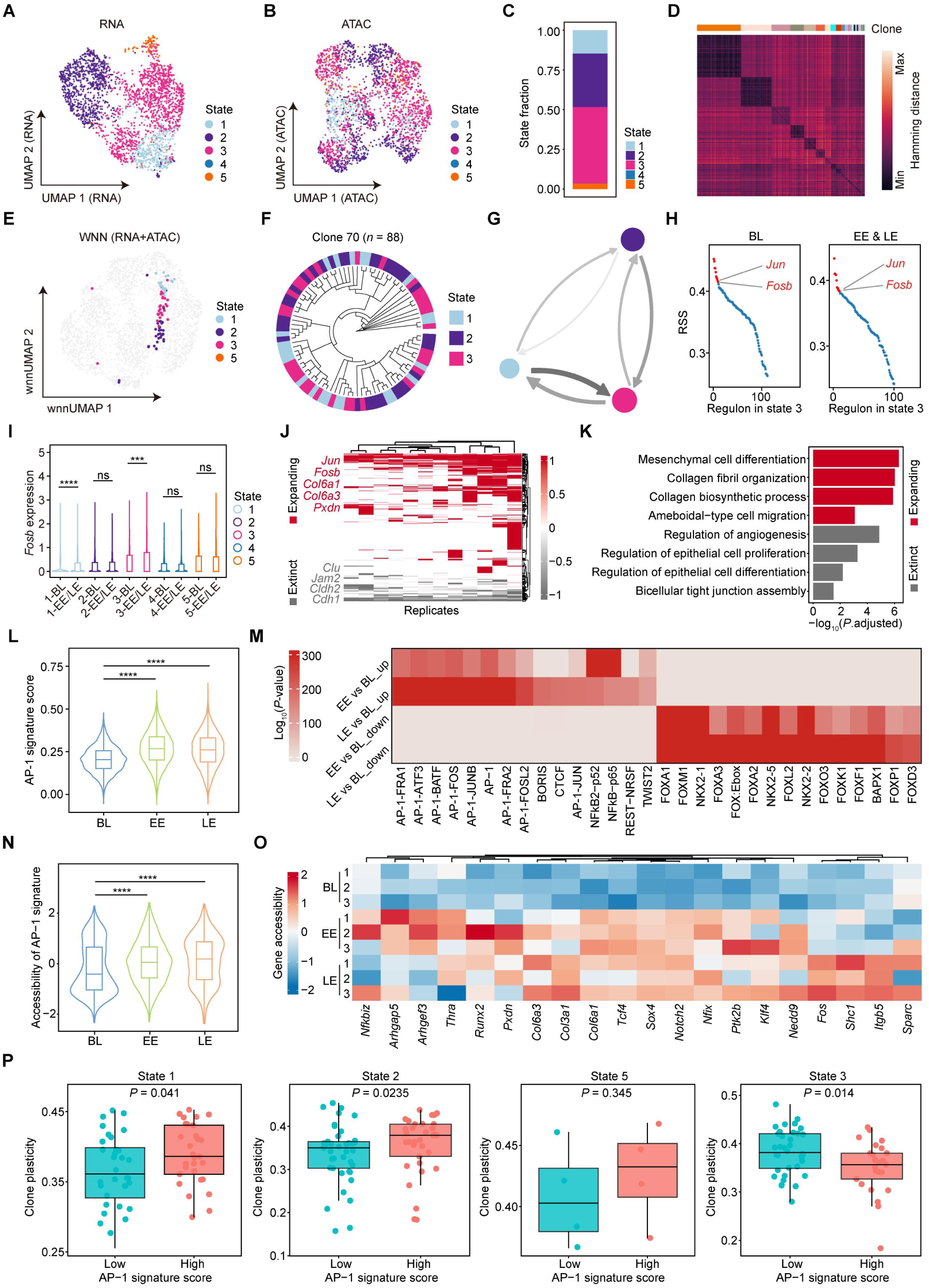
Integrative analysis identifies AP-1 as the key regulator of phenotypic adaptation and stabilization. (A-B) UMAP visualization based on transcriptome (A) and chromatin accessibility (B) data. (C) Stacked bar chart showing the cell state fractions based on scMultiome analysis. (D) Heatmap showing edit distances of evolving barcodes in clones identified through scMultiome analyses. (E-G) The wnnUMAP visualization (E), phylogeny (F), and state-transition network (G) of representative clone 70 based on scMultiome analyses. Arrows indicated the direction of transition and the thickness/color intensity of arrows indicated the transition probabilities (G). Circle size represents proportion of each state in clone 70 (G). (H) The RSS of TFs in state 3 at the BL, EE and LE stages based on scRNA-seq data analyses. The top 10 TFs in red. (I) Statistical analysis of *Fosb* expression across tumor cell states at BL and EE/LE stages in scRNA-seq data. Wilcoxon test. (J) Heatmap displaying differentially expressed genes (DEGs) between expanding clones and extinct clones in scRNA-seq data. Representative DEGs are shown. Data represent biological replicates. (K) Pathway enrichment analysis was performed on DEGs from (J). Representative pathways are shown. Wilcoxon test. (L) AP-1 signature scores in the state 3 across the three stages. Wilcoxon test. (M) Transcription factor enrichment analysis of motifs differentially active across the three stages. Top 15 transcription factors associated with up- and downregulated motif activity are shown. Colors represent enrichment *P-*values. (N) Motif accessibility of AP-1 signature across the three stages. Wilcoxon test. (O) Motif accessibility of representative genes from the AP-1 signature across the three stages. Data represent three technical replicates per stage. (P) Correlation between clonal-level plasticity of each cell state and AP-1 signature score across the three stages. One-sided Wilcoxon test, \**P* < 0.05; \*\**P* < 0.01; \*\*\**P* < 0.001; \*\*\*\**P* < 0.0001; ns, not significant, *P* ≥ 0.05.

**Figure S18.**
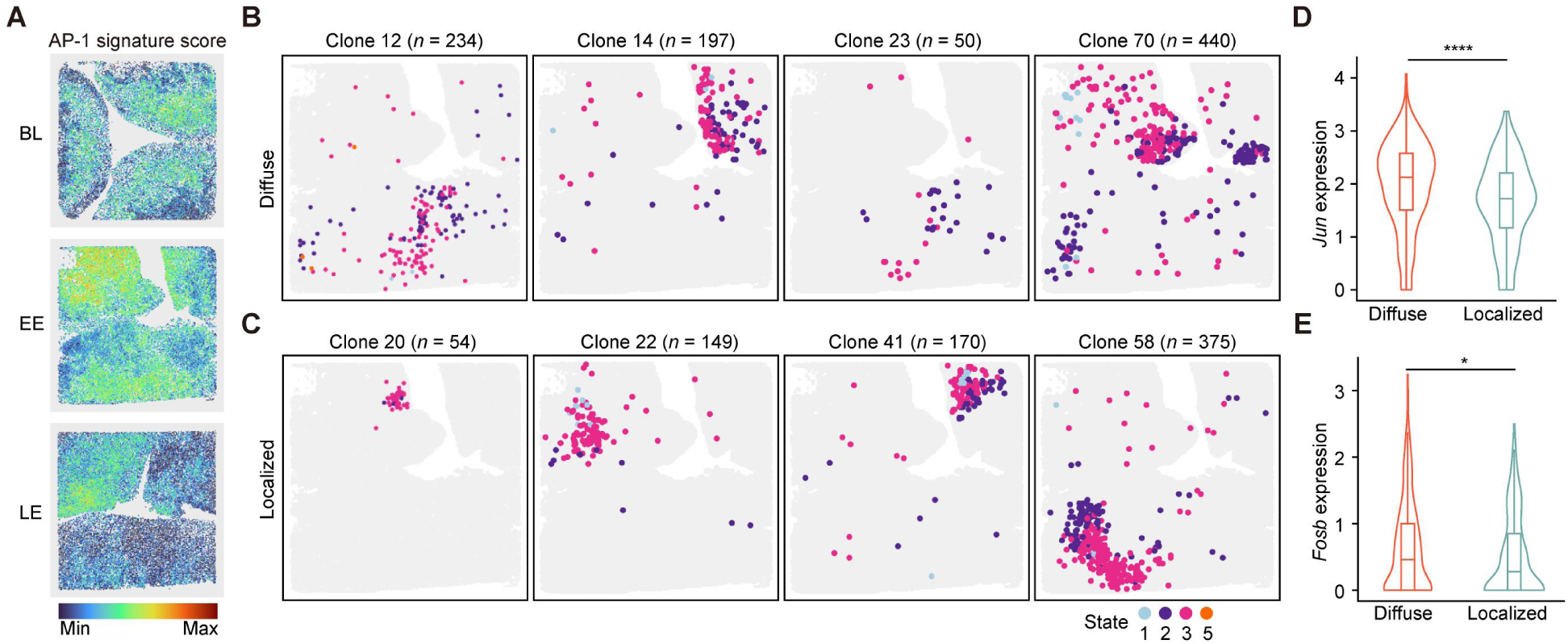
Associations of clonal spatial distribution and AP-1 expression. (A) AP-1 signature score across the three stages through spatial transcriptomic data analyses. (B-C) Spatial mapping of representative clones with relatively diffuse (B) or localized (C) distributions at the EE stage. (D-E) Expression levels of *Jun* (D) and *Fosb* (E) in state 3 were compared between spatially diffuse clones and spatially localized clones at the EE stage based on scRNA-seq data analyses. Wilcoxon test.

**Figure S19.**
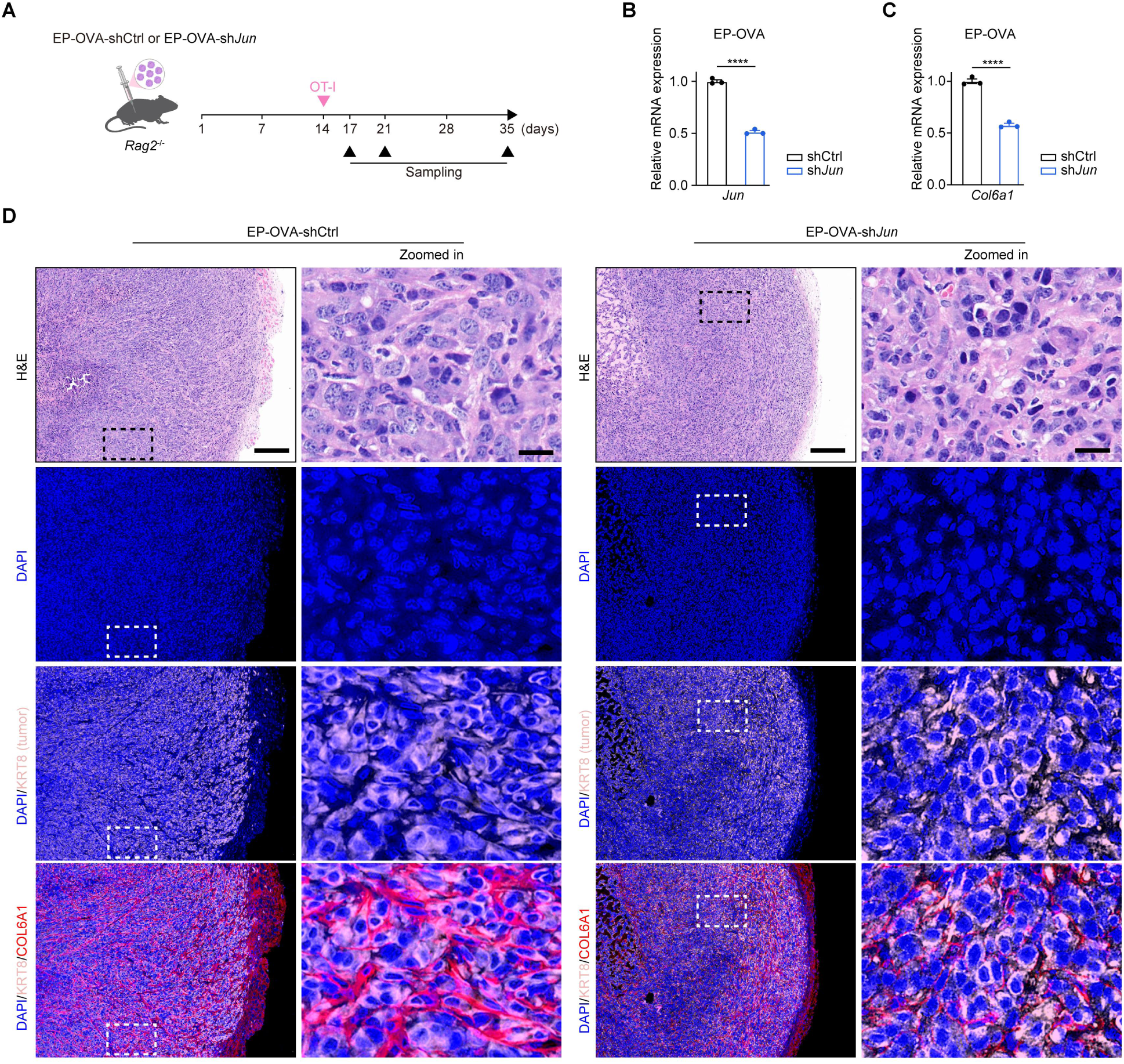
Functional validation of AP-1-mediated immune evasion. (A) Schematic illustration of AP-1 functional validation experiments. 50,000 EP-OVA-shCtrl or EP-OVA-sh*Jun* cells were subcutaneously transplanted into *Rag2*^-/-^ mice. At two weeks post-engraftment, 1 × 10^6^ OT-I cells were injected intravenously, and tumor volumes were monitored approximately every three days. EP-OVA-shCtrl and EP-OVA-sh*Jun* tumors were collected at 3, 7, and 21 days post-OT-I transfer for subsequent multiplex immunofluorescence analyses. (B-C) The expression levels of *Jun* (B) and the representative target gene *Col6a1* (C) in EP-OVA-shCtrl and EP-OVA-sh*Jun* cells were detected by qRT-PCR. Two-sided Student’s t-test. (D) Representative H&E and immunofluorescence images showing COL6A1 expression in tumor samples at day 3 post-OT-I injection. Scale bars represent 200 µm and 20 µm (zoomed-in view), respectively.

